# Identifying Novel Estrogenic Mitochondrial Targets in Hypothalamic Proopiomelanocortin Neurons by Chemoproteomics

**DOI:** 10.64898/2026.06.17.732817

**Authors:** Jian Qiu, Martha A. Bosch, Mya Wolfe, Ksenija Korac, Stefano Rizzo, Todd L. Stincic, Scotland E. Farley, Wendy Fitzgerald, Megha Rajendran, Aurélien Laguerre, Frank Stein, Philip F. Copenhaver, Oline K. Rønnekleiv, Sean M. Rønnekleiv-Kelly, Tatiana K. Rostovtseva, Sergey M. Bezrukov, Carsten Schultz, Martin J. Kelly

## Abstract

Loss of estrogens at menopause is linked to impaired brain metabolism and increased risk of Alzheimer’s disease (AD). However, estrogen replacement therapies are limited due to the deleterious effects of estrogen on peripheral organs and an increased risk of vascular dementia. We have developed a non-steroidal estrogenic compound, STX, which does not bind to the classical estrogen receptors α and β, but mimics estrogenic signaling in the central nervous system (CNS) without eliciting the peripheral reproductive actions. STX is protective against neurodegeneration in stroke and AD models, but its molecular targets are unknown. Here, we identified and validated neural targets of STX using chemoproteomic, molecular biological, electrophysiological and metabolic assays in hypothalamic proopiomelanocortin (POMC) neurons. Chemoproteomic profiling identified voltage-dependent anion channels (VDAC1-3) as major intracellular binding partners in mHypo43 (POMC) cells. Based on quantitative single-cell PCR, *Vdac2* was identified as the dominant isoform in female hypothalamic POMC neurons. Seahorse metabolic flux analyses showed that STX potently increased glycolysis, oxidative respiration and mitochondrial ATP production in mHypo43 cells. Nanomolar concentrations of STX enhanced VDAC2 voltage-dependent gating in reconstituted lipid membranes and shifted the low-conductance states toward anion selectivity, consistent with increased ATP flux. Together, these findings identify VDAC proteins as previously unrecognized mitochondrial targets of STX and demonstrate that STX can directly modulate VDAC2 channel properties and cellular bioenergetics. These findings establish a STX–VDAC mitochondrial mechanism in an estrogen-responsive neuronal system and provide a molecular framework for future studies examining how this pathway contributes to the neuroprotective actions of STX in neurodegenerative diseases.

## INTRODUCTION

Approximately two-thirds of patients with Alzheimer’s disease (AD) are women, and the loss of the ovarian steroid 17β-estradiol (E2) at menopause is strongly associated with increased vulnerability to neurodegeneration (Nebel et al., 2018). Although E2 replacement is neuroprotective in animal models, including non-human primates (Rapp et al., 2003; Morrison et al., 2006), hormone replacement therapy in post-menopausal women has produced adverse outcomes, including increased risk of coronary heart disease and vascular dementia (Rossouw et al., 2002; Shumaker et al., 2003). These limitations have driven the development of neuroprotective selective estrogen receptor modulators (SERMs), but most act through nuclear estrogen receptors and retain undesirable peripheral effects (Murphy et al., 2003).

In search of a selective neuroprotective SERM, we developed STX, a synthetic diphenyl acrylamide compound that selectively targets the central nervous system (CNS) and mimics the rapid, membrane-initiated estrogen signaling without activating classical estrogen receptors (Qiu et al., 2003; Qiu et al., 2006; Tobias et al., 2006; Roepke et al., 2010; Smith et al., 2014). STX activates G protein–coupled signaling pathways in hypothalamic neurons (Qiu et al., 2003; Qiu et al., 2006; Smith et al., 2013). In proopiomelanocortin (POMC) neurons, STX induces heterologous desensitization of inhibitory Gα_i/o_-coupled receptors, including μ-opioid and GABA_B_ receptors, thereby increasing neuronal excitability (Qiu et al., 2003; Qiu et al., 2006). Importantly, STX crosses the blood–brain barrier, can be administered chronically, and reproduces key estrogenic effects on thermoregulation and metabolism without peripheral estrogenic actions (Roepke et al., 2010).

In addition to its rapid neuromodulatory effects, STX exhibits robust neuroprotective properties. It reduces ischemic neuronal damage in hippocampal neurons (Lebesgue et al., 2010), enhances synaptic plasticity, and protects against β-amyloid toxicity in cellular and animal models of AD (Gray et al., 2016; Quinn et al., 2022; Lee et al., 2025). These effects are accompanied by improvements in mitochondrial function, suggesting that regulation of cellular bioenergetics may be a central component of STX action. Mitochondrial dysfunction is a defining feature of neurodegenerative diseases (Ashleigh et al., 2023), and declines in brain metabolic capacity are strongly associated with menopause (Brinton et al., 2015; Mosconi et al., 2018). However, the molecular targets through which STX modulates mitochondrial function and neuronal activity remain unknown.

POMC neurons provide a particularly useful neuronal population in which to investigate estrogen-sensitive mitochondrial mechanisms because they are metabolically responsive, are well-established targets of E2, and represent a neuronal population in which the membrane-initiated actions of STX have been extensively characterized. Moreover, increasing evidence indicates that AD involves hypothalamic dysfunction and metabolic abnormalities in addition to its well-characterized cortical and hippocampal pathology (Ishii and Iadecola, 2015).

Consistent with this concept, Do and colleagues demonstrated hypothalamic neurodegeneration in the 3xTg-AD mouse model, including metabolic abnormalities, increased hypothalamic inflammatory and apoptotic markers, and reductions in arcuate hypothalamic neuronal populations, including POMC-expressing neurons. Voluntary exercise attenuated hypothalamic apoptosis and restored POMC neuronal populations, suggesting that hypothalamic POMC neurons represent a vulnerable and modifiable neuronal population in an experimental AD model (Do et al., 2018). Independent evidence also links POMC-derived signaling to AD-associated neuronal dysfunction. In APP/PS1 mice, hippocampal POMC/MC4R signaling is impaired during the development of synaptic dysfunction, whereas restoration of this pathway improves long-term potentiation, synaptic transmission, and dendritic spine abnormalities (Shen et al., 2016). Together, these observations provide a biological framework for investigating POMC-related mechanisms in the context of neurodegeneration.

However, identifying the molecular targets of small molecules such as STX in intact cells remains a major challenge. Unlike protein–protein interactions, small molecule–protein interactions are often transient and difficult to capture biochemically. Diazirine-based photoaffinity labeling, combined with biorthogonal click chemistry, enables covalent capture and enrichment of ligand-bound proteins in living systems (Moellering and Cravatt, 2012; Schultz et al., 2022). This “flash-and-click” strategy has proven effective for mapping lipid and small-molecule interactomes in complex cellular environments (Haberkant et al., 2013; Hulce et al., 2013; Höglinger et al., 2017; Müller et al., 2020; Müller et al., 2021).

Therefore, we combined chemoproteomics, molecular biology, electrophysiology and mitochondrial physiology to identify the cellular targets of STX in hypothalamic POMC neurons. Using a photoactivatable STX probe, we identified all three VDAC isoforms as prominent intracellular binding partners. We show that STX directly modulates mitochondrial VDAC channels, enhancing voltage-dependent gating and altering ion selectivity, with corresponding increases in mitochondrial ATP production, respiratory capacity and glycolytic function. The present study is therefore primarily a target-identification and mechanistic investigation, and identification of the STX–VDAC interaction provides a molecular mechanism that can be tested in future disease-specific models.

## RESULTS

### Chemoproteomic identification of STX targets reveals VDACs as prominent binding partners

A key regulator of mitochondrial metabolism is the voltage-dependent anion channel (VDAC), located in the outer mitochondrial membrane. VDACs control the exchange of ATP, ADP, and other metabolites between mitochondria and the cytosol, thereby governing cellular energy homeostasis (Rostovtseva and Colombini, 1996; Colombini, 2004; Rostovtseva and Bezrukov, 2008). Modulation of VDAC gating and ion selectivity can directly influence mitochondrial respiration and metabolic flux. Although steroidal compounds have been reported to bind VDACs, their functional effects on channel activity remain incompletely characterized (Camara et al., 2017; Reina and De Pinto, 2017; Rostovtseva et al., 2020). In isolated liver mitochondria, the non-steroidal SERM tamoxifen has been shown to bind VDACs and potentially regulate anion transport across the mitochondrial membrane (Unten et al., 2022). Given the structural similarities between STX and tamoxifen, we therefore asked whether VDACs might also represent molecular targets of STX.

To test this possibility, we designed a novel bifunctional STX derivative (BF-STX) containing both a photo-cross-linkable diazirine group and an alkyne handle, which was synthesized in collaboration with SiChem GmbH (Germany) (**Figure 1A**). BF-STX retained full biological activity, reproducing the ability of STX to induce heterologous desensitization of Gi/o-coupled receptor signaling in POMC neurons, which inhibits the activity of these anorexigenic neurons (Qiu et al., 2003; Qiu et al., 2006) (**Figure 1B**). Also, upon UV activation, BF-STX enabled robust and cell-type–specific labeling in hypothalamic slices from female C57BL/6 mice and mHypo43 cells (**Figures 1C,D**).

**Figure 1.**
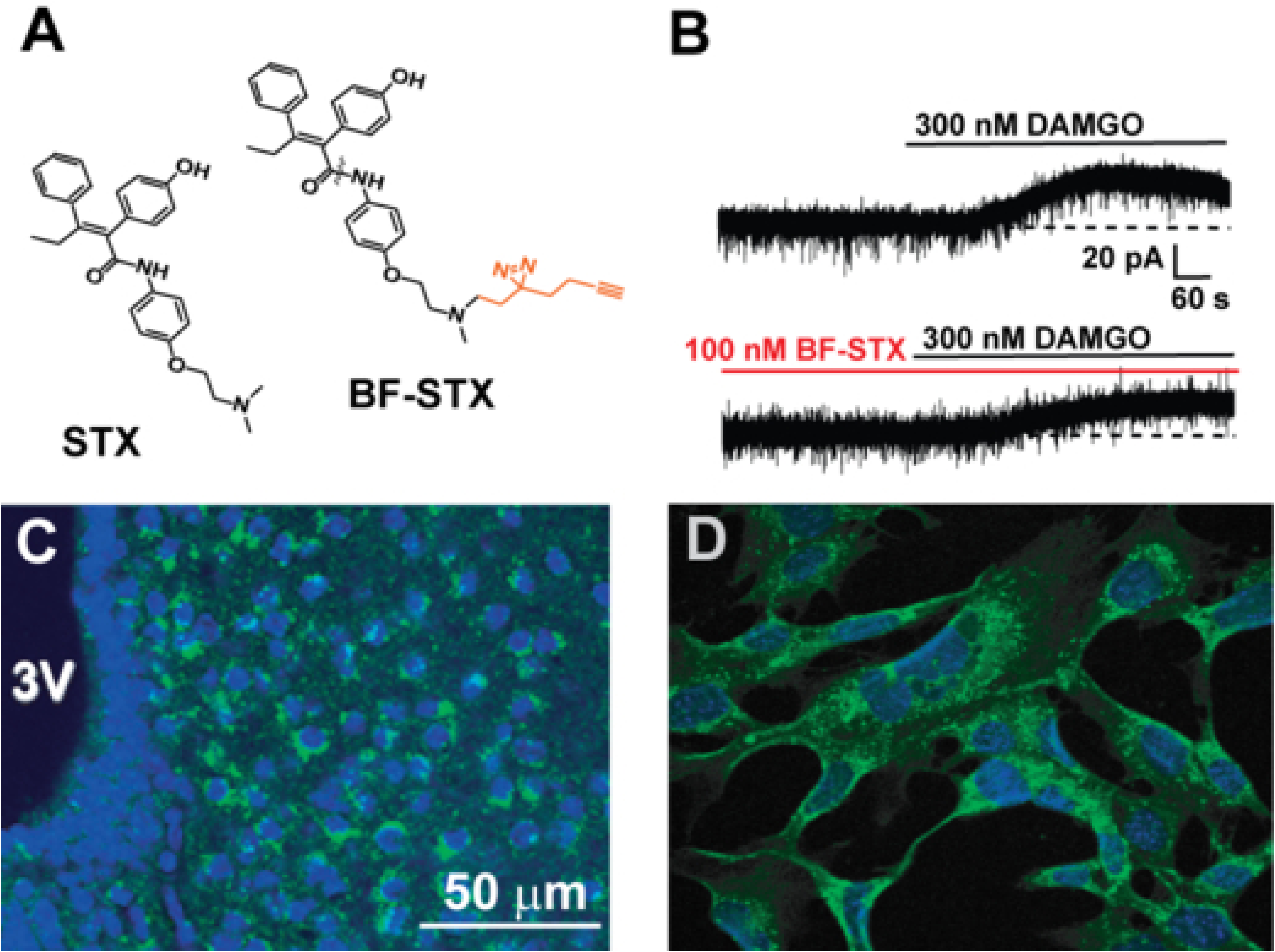
Design, stability, and functional characterization of photoreactive bifunctional-STX (BF-STX) in neurons and cell lines. **A.** Chemical structures of STX and bifunctional-STX (BF-STX), featuring a diazirine group connected to an alkyne handle (both shown in red) for photo-crosslinking and click chemistry, respectively. Upon 7 minutes under 350 nm UV illumination in methanol (MeOH), BF-STX exhibited characteristic changes in proton resonances near the diazirine moiety, confirming photoreactivity (see Supplemental Files). BF-STX remained chemically stable in the dark. **B.** BF-STX rapidly attenuated µ-opioid receptor–mediated responses in POMC^EGFP^ neurons. The µ-opioid receptor, like the GABA_B_ receptor, is G_i/o_-coupled and activates G protein-coupled, inwardly rectifying K^+^ (GIRK) channels in POMC^EGFP^ neurons. In voltage-clamp recordings (V_hold_ = -60 mV), an EC₅₀ concentration of DAMGO (a µ-opioid receptor agonist) elicited GIRK-mediated outward currents (top trace), which were reduced by 38.7 ± 5.7 % (n=3) following a 15-minute exposure to BF-STX. This heterologous desensitization is consistent with previous observations using the BF-STX parent compound, STX, which causes a 41% attenuation in the GABA_B_ (and µ-opioid) receptor-mediated activation of GIRK channels in POMC neurons (Qiu et al., 2003). **C,D.** Confocal images showing subcellular accumulation of BF-STX (10 µM, 30 min). BF-STX treated cells were subjected to UV crosslinking (2.5 min at 350 nm, on ice), cell fixation, and fluorescent labelling via copper-click reaction using picolyl-azide Alexa Fluor 488. **C.** In hypothalamic slices, BF-STX selectively labeled POMC neurons near the third ventricle (3V). **D.** BF-STX labeled mHypo43 cells.

To identify candidate STX-binding proteins, mHypo43 POMC cells were incubated with BF-STX (10 μM) for 30 min and then exposed to UV light (350 nm) to induce crosslinking (**Figure 2A**). Negative controls included samples without UV exposure and cells incubated with BF-STX together with a 3-fold higher concentration of STX. These controls ensured that the enriched hits were legitimate targets and not “off-targets” from photo-crosslinking (Kleiner et al., 2017; West et al., 2021). Protein–BF-STX conjugates were labeled via click chemistry using picolyl-azide agarose beads, digested on-bead, TMT-labeled, and analyzed by LC-MS/MS on an Orbitrap Fusion Lumos system at EMBL as described in the Materials and Methods. VDAC1, VDAC2, and VDAC3 emerged as prominent targets (**Figure 2A**, **Suppl Table 1**). Interestingly, even with a short exposure (5 min) to BF-STX, VDACs emerged as significant targets (**Figure 2B, Suppl Table 2**), which is congruent with the rapid neuroprotective actions of STX to rescue CA1 hippocampal neurons following global ischemia in female rats (Lebesgue et al., 2010). Furthermore, fold-change correlation plots showed that VDAC proteins were strongly enriched in the UV crosslinking condition relative to the no-UV control (x-axis), while enrichment was reduced in the competition experiments **(Figure S4A**). Competition with unlabeled STX reduced BF-STX labeling of several candidate proteins (**Figure S4B**). Among the VDAC isoforms, the most pronounced competition was observed for VDAC3, with the VDAC3 signal no longer detected at three molar equivalents of unlabeled STX. The low-abundance VDAC1 signal remained detectable under these competition conditions, whereas VDAC2 labeling was reduced but not eliminated. However, these proteomic data are not suitable to provide quantitative measurements of competition or relative binding affinities among the VDAC isoforms. Thus, these results support VDAC proteins as candidate STX-interacting proteins but do not establish differences in STX binding affinity or specificity among VDAC1–3.

**Figure 2.**
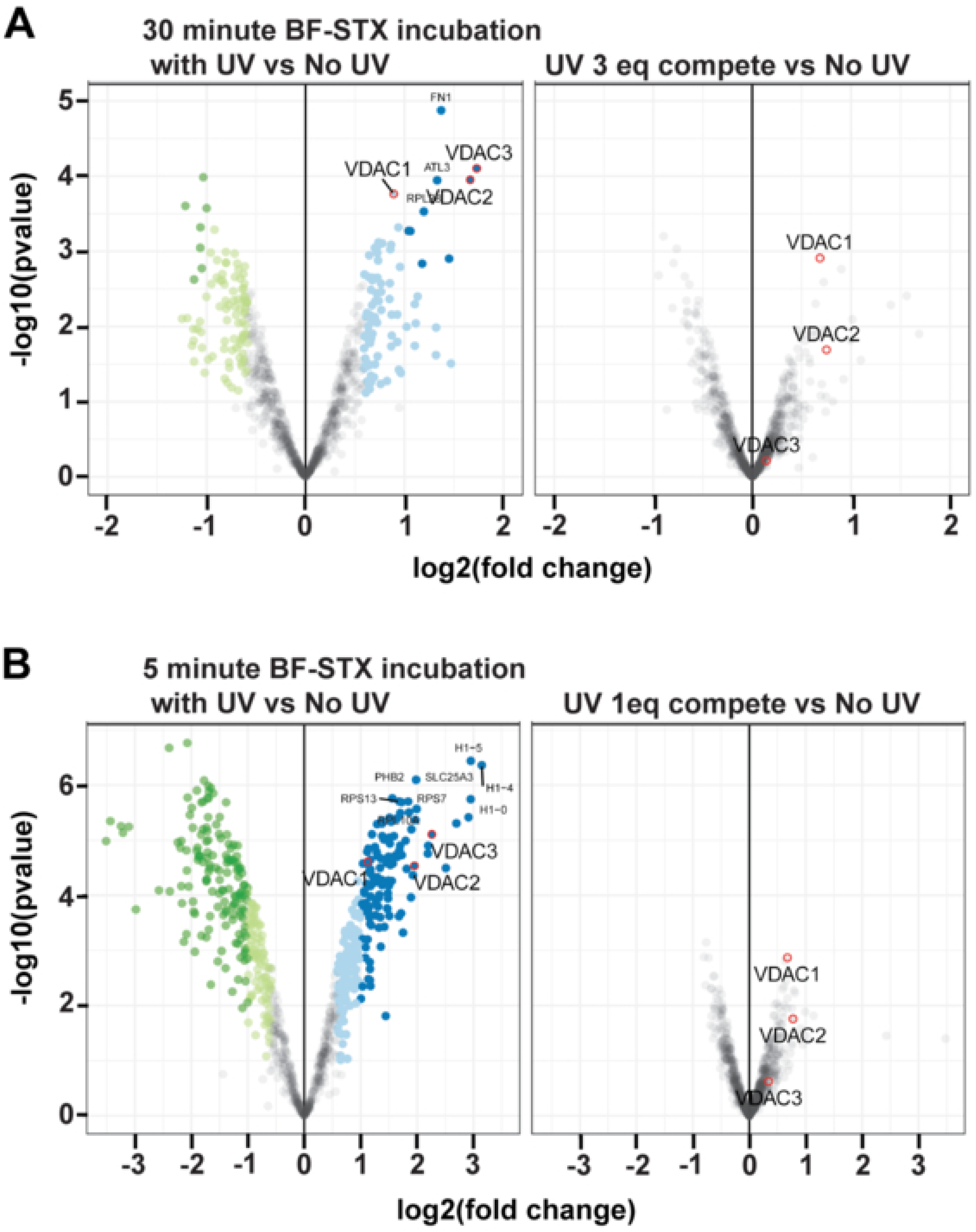
Chemoproteomic identification of BF-STX–interacting proteins in mHypo43 cells following UV crosslinking. **A**. Volcano plots showing enriched proteins identified after 30 min BF-STX incubation under UV crosslinking conditions compared with no-UV controls (left), and after competition with 3× excess unlabeled STX during UV crosslinking compared with no-UV controls (right). Proteins significantly enriched by BF-STX labeling are highlighted, with VDAC1, VDAC2, and VDAC3 among the prominent candidate targets identified after UV exposure. Competition with excess STX reduced enrichment of VDAC family proteins, consistent with competition-sensitive BF-STX labeling. **B.** Volcano plots showing enriched proteins identified after 5 min BF-STX incubation under UV crosslinking conditions compared with no-UV controls (left), and competition with 1× excess unlabeled STX during UV crosslinking compared with no-UV controls (right). Shorter BF-STX incubation increased enrichment and statistical significance of VDAC1, VDAC2, and VDAC3. Competitive STX treatment reduced enrichment of these proteins, consistent with BF-STX associated labeling of VDAC family members. The x-axis represents log2 (fold change), and the y-axis represents –log10 (p value).

### VDAC2 is the predominant isoform expressed in POMC neurons

At the molecular level, we first examined whether VDACs are expressed in hypothalamic anorexigenic POMC neurons, given their central role in mitochondrial metabolite exchange. Single-cell qPCR from pooled POMC neurons from female mice revealed a clear expression hierarchy, with *Vdac2 > Vdac3 >> Vdac1* mRNA (**Figures 3A,B**), indicating that Vdac2 was the predominant transcript among the three isoforms in the POMC neurons examined. This distribution contrasts with most tissues, where VDAC1 predominates (Zinghirino et al., 2021), suggesting a cell-type–specific specialization of mitochondrial outer membrane channels in POMC neurons.

**Figure 3.**
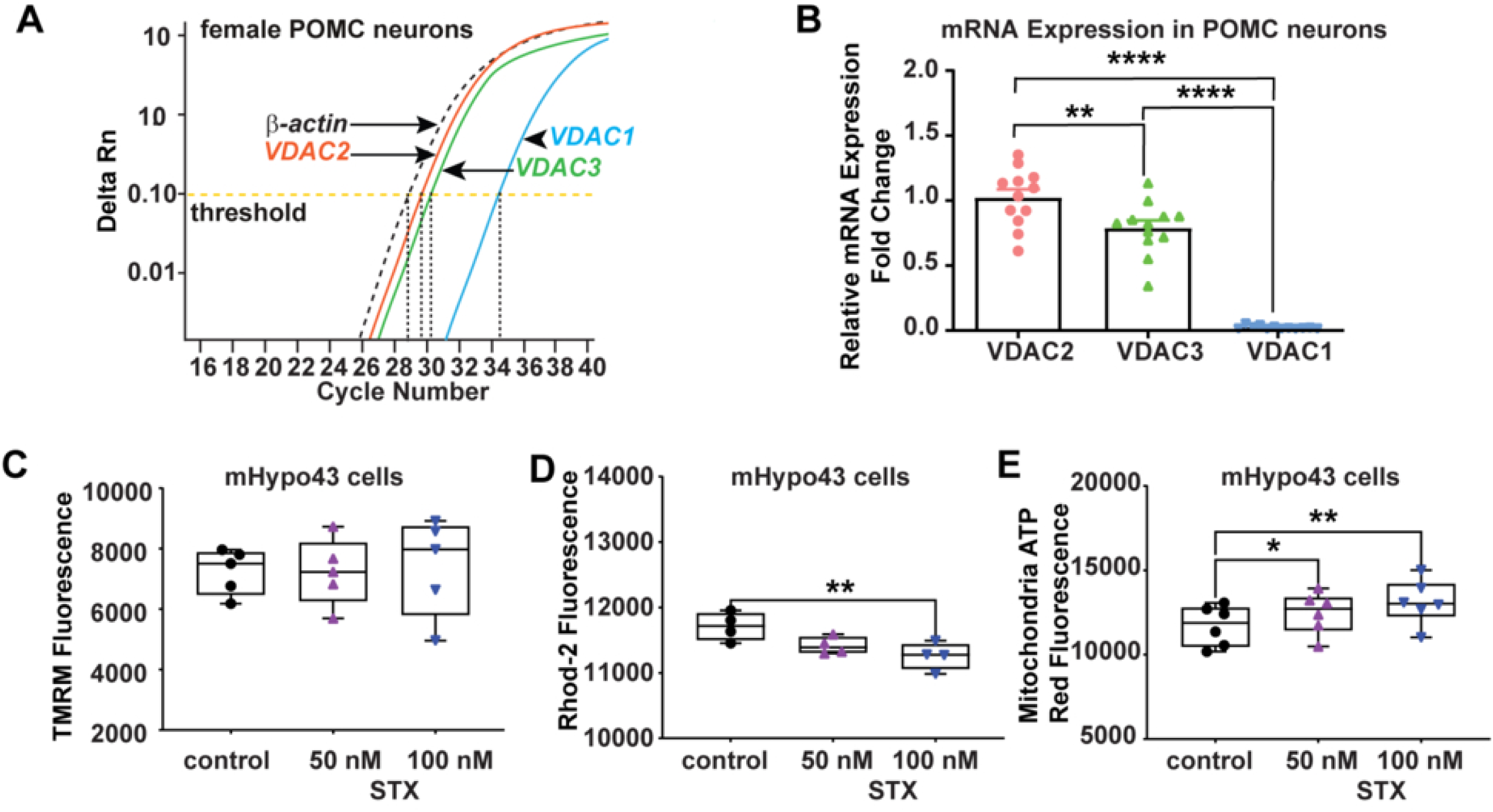
qPCR identification of *Vdac1-3* mRNA expression in POMC neurons. **A.** Quantitative PCR (qPCR) amplification curves for VDAC1-3 were generated from 10-cell pools of *Pomc^EGFP^* neurons in female mice. Cycle number was plotted against normalized fluorescence intensity (ΔRn) to visualize amplification, with the cycle threshold (Ct; dashed line) indicating the point at which fluorescence exceeded background levels and sample values were quantified. **B**. Summary bar graphs of the relative expression of *Vdac2, Vdac3, Vdac1* mRNA in female POMC neurons (12 ten-cell pools from three animals/group). Statistical analysis was performed using one-way ANOVA followed by Tukey’s post hoc test (****p<0.001, **p<0.01). **C**. Mitochondrial membrane potential determination using TMRM dye; **D.** Mitochondrial calcium using Rhod2 dye; and **E,** mitochondrial ATP using BioTracker ATP-Red live cell dye were measured in mHypo43 cells upon treatment with 50 and 100 nM STX. Data from 4-6 independent experiments are represented in the box plots. The symbols represent data from each repeat, the borders of the boxes define the 25^th^ and 75^th^ percentiles, with the median displayed as black lines, and error bars indicate the standard deviation from the mean. Statistical analysis was performed using one-way ANOVA followed by Dunnett’s *post hoc* test ( **p<0.01, *p<0.05).

To assess whether STX influences mitochondrial function in these cells, we measured mitochondrial membrane potential, calcium and ATP levels in mHypo43 neurons. STX (100 nM) did not alter the mitochondrial membrane potential, but it significantly reduced mitochondrial calcium concentrations and increased mitochondrial ATP levels (**Figures 3C-E**). These findings indicate that STX enhances mitochondrial function without inducing depolarization, consistent with improved metabolic efficiency rather than mitochondrial stress.

### STX directly modulates VDAC2 channel gating and ion selectivity

To determine whether STX directly targets VDAC channels, we performed electrophysiological recordings of recombinant human VDAC2 reconstituted in planar lipid membranes (**Figure 4A**) (Rostovtseva and Bezrukov, 2015). VDAC2 was selected because it showed the highest mRNA expression in POMC neurons (**Figure 3A,B**) and, unlike VDAC3, can be more reliably reconstituted into artificial membranes (Queralt-Martín et al., 2020). This focus does not exclude other VDAC isoforms as STX targets, as our chemoproteomic experiments identified all three VDACs, with particularly strong competition of VDAC3 labeling by unlabeled STX. In a symmetrical 1 M KCl buffer solution, VDAC2 typically forms channels of ∼3.5 – 4.0 nS conductance (**Figure 4A**) (Rosencrans et al., 2025), which characteristically transition (*i.e*., gate) from the high conducting “open” state to a variety of lower-conducting or “closed” states in response to large applied voltages (**Figure 4B**, control trace). Under these experimental conditions, the application of 60 mV was required to observe consistent voltage gating. For the sake of illustrative clarity, in **Figure 4B** we show traces at negative potentials. The addition of 100 and 300 nM STX to the “cis” (grounded) side (**Figure 4A**) of the membrane caused channel closure at lower negative potentials, -20 and -10 mV, respectively (**Figure 4B**, lower traces). The transitions to the low conductance states were reversible such that the voltage jump to 0 mV fully reopened the channel, a characteristic behavior of reconstituted VDACs (Colombini, 1989). The single-channel recordings revealed that in the presence of STX, voltages of 40-50 mV lower than those in the control were sufficient for VDAC2 single-channel gating.

**Figure 4.**
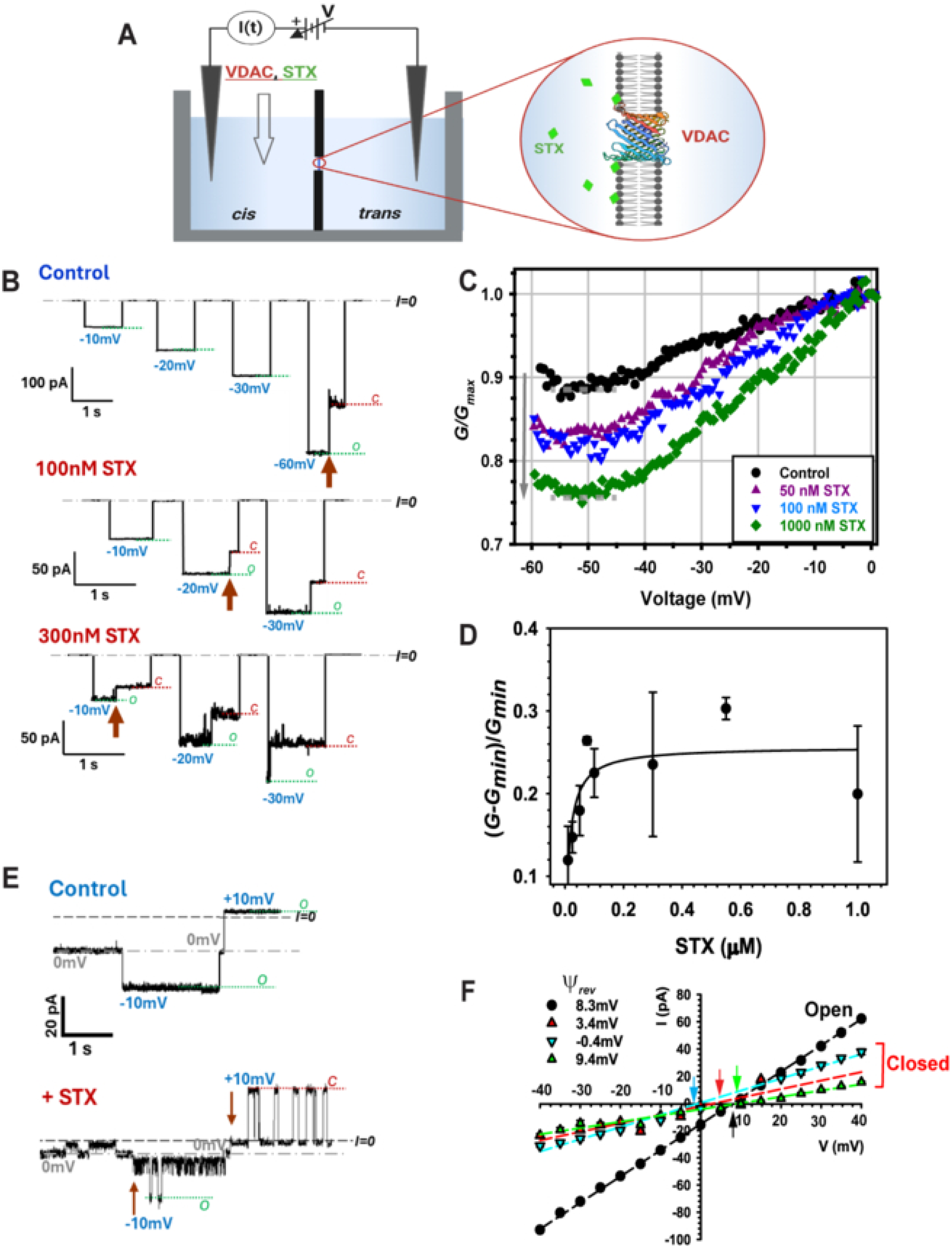
STX promotes voltage gating of VDAC2 and shifts the channel’s low-conductance states toward anion selectivity. **A.** A schematic of the experimental setup for VDAC reconstitution. In these experiments, a single VDAC spans a planar lipid membrane that separates and electrically isolates two buffer-filled compartments. The ionic current through the channel is generated by the applied voltage and constantly recorded. VDAC and STX (green diamonds) were added to the cis compartment. **B.** Electrophysiological recordings of the recombinant human VDAC2 reconstituted into planar lipid membranes before (Control) and after the addition of 100 and 300 nM STX to the membrane-bathing solution. The current traces, representing two VDAC2 channels, were recorded on the same membrane at the indicated voltages, which were returned to 0 mV between each subsequent voltage application. Characteristic voltage-gating behavior is seen as a stepwise current transition from the unique high-conducting or “open” state (“o” state, indicated by green dashed lines) to a low-conducting “closed” state (“c” state, indicated by red dashed lines). The lowest voltages at which gating was observed at each condition are indicated by red arrows. Here and in panel E, the dash-dotted gray lines show the zero current level (I=0). Membrane bathing solutions contained 1 M KCl buffered with 5 mM HEPES at pH 7.4. **C.** The normalized VDAC2 conductance as a function of the applied voltage obtained in a representative experiment with about 60 VDAC2 channels reconstituted into a planar membrane without (Control) and with subsequent additions of 50, 100, and 1000 nM of STX. Normalized conductance is defined as G/G_max_, where G is the average conductance at voltage V, and G_max_ is the maximum average conductance at voltages close to 0 mV. Gating behavior was assessed by applying a triangular voltage wave of ±60 mV, 5 mHz. The addition of STX resulted in a decrease in the minimal (G_min_) normalized conductance (indicated by dashed gray lines) at the application of negative voltages (emphasized by the downward arrow), indicative of increased gating. These results, obtained in multichannel membranes, are consistent with the single-channel records in B. Other experimental conditions were as in (B). **D**. STX promotes VDAC2 voltage gating in a dose-dependent manner. Normalized gating response of four independent experiments, as in (C), represented as (G_max_-G_min_)/G_min_ and plotted against increasing concentrations of STX. Each data point denotes the mean ± SD (n=4). The line is a ligand-binding, one-site saturation curve with K_d_ = 15 ± 4 nM. **E**,**F** STX affects the ion selectivity of VDAC2 voltage-induced “closed” states. **E.** Representative single-channel trace obtained in 1 M (cis)/ 200 mM (trans) KCl gradient before (Control, upper trace) and after (+ STX, lower trace) addition of 100 nM STX to the cis compartment at the indicated voltages. Dashed green and red lines indicate open and “closed” states, respectively; dashed gray lines indicate zero current. A planar lipid membrane was formed from DPhPC. **F.** I/V curves obtained from the traces, examples of which are shown in E, for the open (black circles) and three closed (triangles) states. Linear regressions (dashed lines) allow calculation of the reversal potential (ψ_rev_, indicated by arrows) of each state (shown in the inset). A positive ψ_rev_ corresponds to anion selectivity, whereas a negative ψ_rev_ corresponds to cation selectivity. The open state with conductance of 1.9 nS is anion selective (ψ_rev_ = 8.3 mV); two low-conducting states of 0.5 and 0.6 nS are also anion selective with ψ_rev_ equal to 9.4 and 3.4 mV, respectively, and the low-conducting state of 0.9 nS conductance, is non-selective (ψ_rev_ = 0.4 mV).

A quantitative analysis of the reconstituted VDAC voltage gating requires recording from many channels, which is best achieved using a multichannel approach (Rostovtseva et al., 2006; Teijido et al., 2014; Rappaport et al., 2015). In this protocol, slow triangular voltage waves (5 mHz, ±60 mV) are applied to the membrane containing 20-100 channels. **Figure 4C** shows representative normalized conductance-voltage plots of G/G_max_ versus voltage at negative potentials, where G is the conductance at a given voltage, and G_max_ is the maximum conductance at voltages close to 0 mV, in control and after addition of 50, 100, and 1000 nM STX to the cis compartment. The STX-enhanced voltage gating was manifested as a decrease in the minimal conductance (G_min_, indicated by dashed gray lines) calculated at |V| > between 45-55 mV (downward arrow in **Figure 4C**). The dose-dependence of STX-enhanced VDAC2 voltage gating was quantified as the normalized change in conductance (G_max_-G_min_)/G_min_ plotted against STX concentrations (**Figure 4D**). The averaged normalized reduction in G_min_ with STX concentration was approximated by a first-order binding curve with a K_d_ of 15 ± 4 nM.

These results suggest a direct functional interaction between STX and VDAC2. To further test this hypothesis, we measured the ionic selectivity of VDAC2 in the presence of STX. The open state of VDAC2, similarly to VDAC1 and VDAC3 (Tan and Colombini, 2007; Queralt-Martín et al., 2020; Rosencrans et al., 2025), is anion-selective, and multiple voltage-induced closed states are characterized by a wide range of low conductances and predominantly cationic selectivity (Colombini, 1989). We measured VDAC2 selectivity in a 5x salt gradient, 1 M KCl (cis side) versus 0.2 M KCl (trans side). A representative trace of VDAC2 obtained in this salt gradient before and after the addition of 100 nM STX to the cis compartment is shown in **Figure 4E**. In the presence of STX, the fluctuations of channel conductance between different conductance levels were measured at ±10 mV. The corresponding I/V plots for the open and the three low-conducting states are shown in **Figure 4F**. The linear regressions through the data points determined the substate conductance as a slope, and the reversal potential Ψ_rev_ as the voltage at which the current is zero for each substate. While the selectivity of the open state (1.9 nS) remained unchanged (Ψ_rev_ = 8.3 mV), in the presence of STX two low-conducting states of 0.5 and 0.6 nS displayed anion selectivity with Ψ_rev_ equal to 9.4 and 3.4 mV, respectively, with one low-conducting state (0.9 nS) being non-selective (inset in **Figure 4F**). These results indicate that STX may alter the selectivity of VDAC2 closed states, without affecting either the conductance or selectivity of the open state.

The absence of an effect of STX on VDAC2 open-channel conductance suggested that this hydrophobic compound most likely interacts with VDAC2 at the protein-lipid interface. We further postulate that by shifting the selectivity of VDAC2 closed states toward anion selectivity, STX modulates the transport properties of VDAC2 to favor ATP flux (Rostovtseva and Colombini, 1996). These results also explain the decreased mitochondrial calcium and increased mitochondrial ATP measured in mHypo43 (POMC) neurons (**Figures 3D,E**). Facilitation of ATP/ADP fluxes may contribute to the neuroprotective effects of STX in CNS neurons (Gray et al., 2016; Quinn et al., 2022; Lee et al., 2025).

### STX enhances mitochondrial respiration and glycolytic function

Based on the finding that STX increased ATP concentrations in mHypo43 cells (**Figure 3E**), we decided to measure cellular metabolism in real-time using the Agilent Seahorse Assay system. We investigated the effects of STX on mitochondrial function, including mitochondrial respiration and glycolysis over time. Mitochondrial respiration (**Figures 5, 6**) was assessed by administering agents that target different components of the electron transport chain (i.e., oligomycin, FCCP and rotenone/antimycin A) (Divakaruni et al., 2014) (see Materials and Methods). These measurements were conducted under control conditions (vehicle) and compared to treatment with 1 nM, 5 nM or 10 nM STX in mHypo43 cells. STX treatment led to an increase in several key parameters of mitochondrial respiration (**Figure 5A**). Quantitative analysis revealed that STX significantly increased basal respiration (**Figure 5B**), ATP production (**Figure 5C**), and maximal respiratory capacity (**Figure 5D**). Additionally, STX augmented spare respiratory capacity (**Figure 5E**), suggesting enhanced mitochondrial adaptability under stress conditions. Notably, proton leak (**Figure 5G**) but not non-mitochondrial respiration (**Figure 5F**) was significantly increased. Since neurons are highly dependent on glucose uptake and glycolysis as a source of energy (Li et al., 2023), we also analyzed glycolysis by exposing mHypo43 cells to oligomycin, FCCP and rotenone/antimycin A, while measuring the extracellular acidification rate (ECAR) over time. There were differences in ECAR with STX treatment versus control (**Figure 5H**). The lower concentrations of STX (1 and 5 nM) significantly increased glycolysis (**Figure 5I**), glycolytic capacity (**Figure 5J**), glycolytic reserve (**Figure 5K**) and non-glycolytic acidification (**Figure 5L**). However, because STX exhibited a non-monotonic dose-response, with 10 nM STX failing to increase glycolytic function, we speculated that estrogens present in the serum might saturate the STX targets and therefore causing a desensitization. Therefore, we serum-starved the mHypo43 cells for 6 hours prior to the Seahorse assay in order to remove any endogenous estrogens (**Figure 6**). The serum starvation did not alter the “bell-shaped” dose-response for glycolytic function (**Figures 6H-L**) and even showed a monotonic dose-response for mitochondrial respiratory parameters (**Figures 6A-G**). Since hepatocytes exhibit the highest mitochondrial density within the human body (Xu et al., 2023), we sought to see if STX was equally potent in immortalized human hepatocyte (IHH) cells to increase mitochondrial respiration. Indeed, 1 nM (and 10 nM) STX increased mitochondrial respiration in both serum (**Figures S5A-G**) and serum-starved conditions (**Figures S6A-G**). As one would predict, since IHH cells are not dependent on glucose uptake as an energy source, glycolytic function was essentially unaltered with STX treatment in serum (**Figures 5H-L**) and serum-starved conditions (**Figures 6H-L**). Collectively, these results indicate that STX acts independently of endogenous estrogenic compounds and are consistent with the fact that neurons require glucose uptake and glycolysis to maintain their electrophysiological activity (Li et al., 2023).

**Figure 5.**
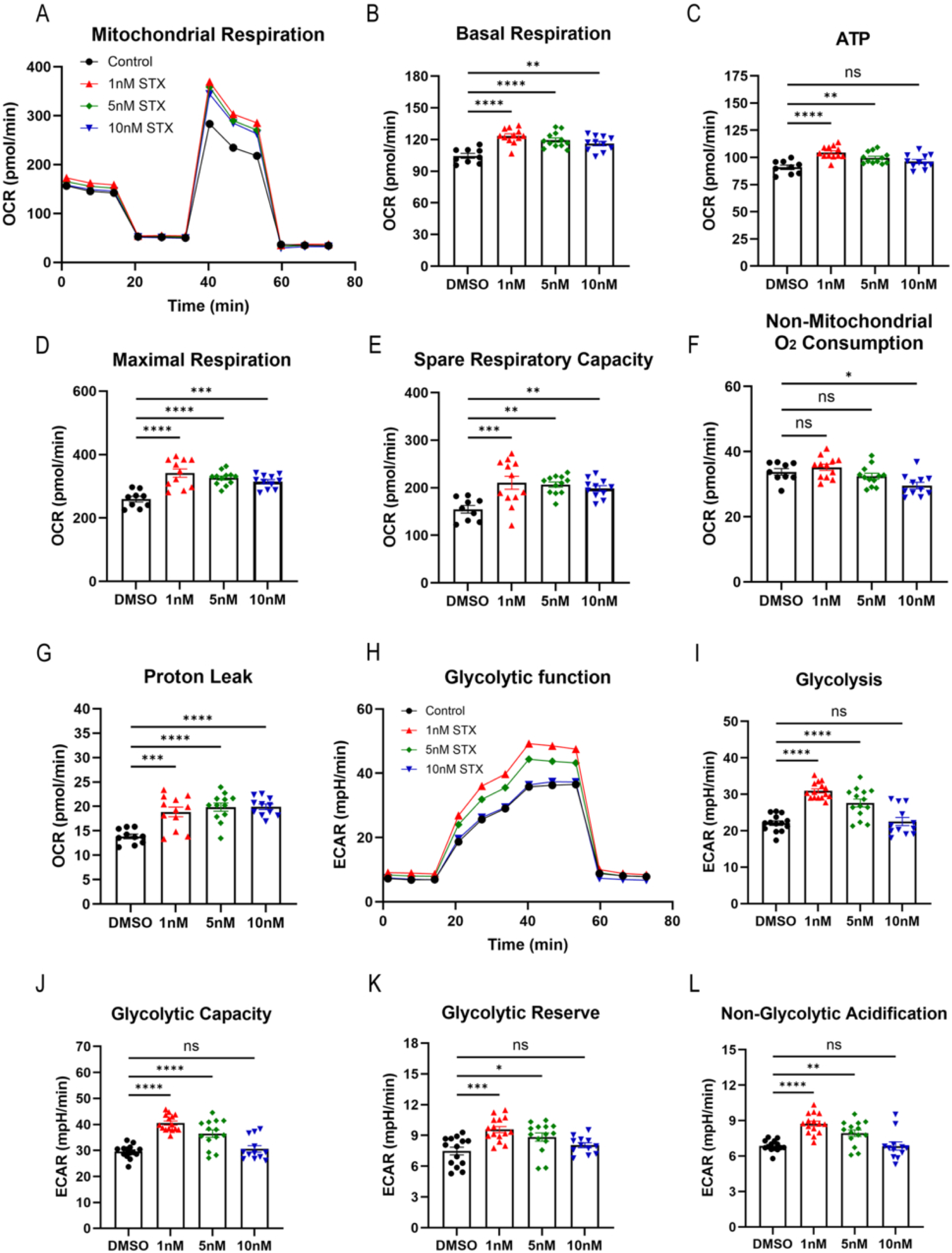
STX induces coordinated mitochondrial and glycolytic activation in non-serum starved mHypo43 cells. **A.** Representative Seahorse XF traces of oxygen consumption rate (OCR) in non-serum starved mHypo43 cells treated with DMSO (control) or STX (1, 5, 10 nM). Sequential injections of oligomycin, FCCP, and rotenone/antimycin A were used to assess mitochondrial respiratory function. **B–G**. Quantification of mitochondrial respiration parameters derived from OCR measurements, including basal respiration (**B**), ATP-linked respiration (**C**), maximal respiration (**D**), spare respiratory capacity (**E**), non-mitochondrial oxygen consumption (**F**), and proton leak (**G**). STX treatment significantly increased basal respiration, ATP-linked respiration, maximal respiration, and spare respiratory capacity, with the most pronounced effects observed at 1 nM. These enhancements were maintained, though slightly attenuated, at 5 nM and 10 nM. Proton leak was significantly increased across STX-treated groups, while non-mitochondrial respiration showed modest changes. **H**. Representative extracellular acidification rate (ECAR) traces during the glycolysis stress test under the same treatment conditions. Sequential injections of glucose, oligomycin, and 2-deoxyglucose (2-DG) were used to assess glycolytic function. **I–L**. Quantification of glycolytic parameters derived from ECAR measurements, including glycolysis (**I**), glycolytic capacity (**J**), glycolytic reserve (**K**), and non-glycolytic acidification (**L**). STX at 1 nM and 5 nM significantly increased glycolysis and glycolytic capacity. At 1 nM, STX significantly increased glycolytic reserve and non-glycolytic acidification, with moderate effects observed at 5 nM. No significant enhancement was observed at 10 nM. Data are presented as mean ± SEM with individual data points shown. Statistical significance was determined by one-way ANOVA followed by Dunnett’s multiple comparison test. ns-not significant, *p < 0.05, **p < 0.01, ***p < 0.001, ****p < 0.0001.

**Figure 6:**
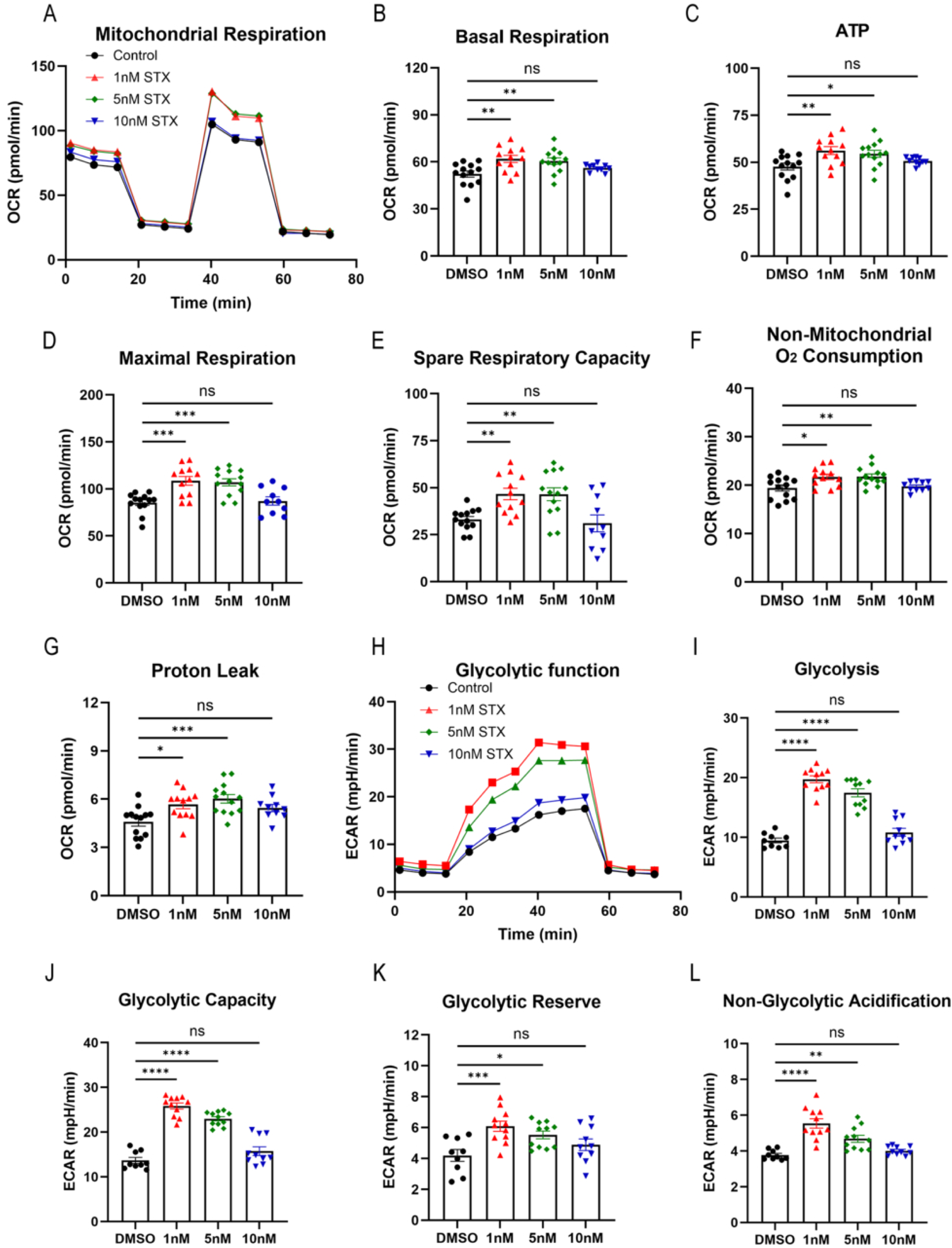
STX induces coordinated mitochondrial and glycolytic activation in serum starved mHypo43 cells. **A**. Representative Seahorse XF traces of oxygen consumption rate (OCR) in 6-hr serum starved mHypo43 cells treated with DMSO (control) or STX (1, 5, 10 nM). Sequential injections of oligomycin, FCCP, and rotenone/antimycin A were used to assess mitochondrial respiratory function. **B–G**. Quantification of mitochondrial respiration parameters derived from OCR measurements, including basal respiration (**B**), ATP-linked respiration (**C**), maximal respiration (**D**), spare respiratory capacity (**E**), non-mitochondrial oxygen consumption (**F**), and proton leak (**G**). STX at 1 nM significantly increased basal respiration, ATP-linked respiration, maximal respiration, and spare respiratory capacity compared to control. Similar but less pronounced effects were observed at 5 nM. In contrast, 10 nM STX showed diminished or non-significant effects across most parameters. Proton leak and non-mitochondrial respiration were modestly increased at lower doses but showed no significant differences at higher concentrations. **H**. Representative extracellular acidification rate (ECAR) traces during the glycolysis stress test under the same treatment conditions. Sequential injections of glucose, oligomycin, and 2-deoxyglucose (2-DG) were used to assess glycolytic function. **I–L**. Quantification of glycolytic parameters derived from ECAR measurements, including glycolysis (I), glycolytic capacity (**J**), glycolytic reserve (**K**), and non-glycolytic acidification (**L**). STX at 1 nM and 5 nM significantly increased glycolysis and glycolytic capacity. At 1 nM, STX significantly increased glycolytic reserve and non- glycolytic acidification, with moderate effects observed at 5 nM. In contrast, 10 nM STX did not significantly enhance these parameters, consistent with a non-monotonic dose–response relationship. Data are presented as mean ± SEM with individual data points shown. Statistical significance was determined by one-way ANOVA followed by Dunnett’s multiple comparison test. ns-not significant, *p < 0.05, **p < 0.01, ***p < 0.001, ****p < 0.0001.

## DISCUSSION

A central strength of this study is the use of photoaffinity labeling coupled with click chemistry to capture STX–protein interactions in intact cellular systems. Because small molecule–protein interactions are often weak, reversible and difficult to preserve during biochemical isolation, the diazirine-containing BF-STX probe provided a means to covalently stabilize ligand-associated proteins after UV activation (Moellering and Cravatt, 2012; Schultz et al., 2022). The alkyne handle then enabled selective enrichment of BF-STX–labeled proteins through copper-catalyzed azide–alkyne cycloaddition, thereby reducing background and increasing confidence in target identification. Importantly, the use of no-UV controls and competition with excess unlabeled STX strengthened the specificity of the chemoproteomic hits and provided complementary evidence that the identified proteins represent STX-interacting candidates rather than nonspecific products of the enrichment procedure.

Our rationale for focusing on POMC neurons derives from their well-established roles in metabolic regulation and estrogen responsiveness, together with the extensive prior characterization of membrane-initiated STX signaling in this neuronal population (Qiu, 2003, 2006; Conde 2107, 20243). Although POMC neurons may not represent a primary or early site of classical AD pathology, they do provide a physiologically relevant and experimentally well-characterized neuronal system in which to investigate the molecular mechanisms underlying STX action. Moreover, AD is increasingly recognized as involving hypothalamic and metabolic dysfunction in addition to its canonical cortical and hippocampal pathology (Ishii and Iadecola, 2015). In the 3xTg-AD mouse model, hypothalamic neurodegeneration is accompanied by a reduction in POMC neuronal populations, an effect that can be attenuated by voluntary exercise (Do et al., 2018). Furthermore, disruption of hippocampal POMC/MC4R signaling has been demonstrated in APP/PS1 mice, whereas restoration of this pathway improves synaptic plasticity and structural abnormalities (Shen et al., 2016). Collectively, these observations provide a broader biological context for examining POMC-associated mechanisms under experimental conditions relevant to AD, while avoiding the implication that POMC neurons constitute a primary locus of AD pathology. Therefore, the enrichment of VDAC1–3 under BF-STX labeling conditions, together with loss of signal after STX competition, supports the conclusion that VDACs are specific intracellular STX-interacting candidates rather than nonspecific contaminants. Thus, the flash-and-click approach provided a mechanistic bridge between the known cellular actions of STX (Lee et al., 2025) and the subsequent functional validation of VDAC2 gating and mitochondrial bioenergetics.

Given the abundant evidence that mitochondrial dysfunction plays a role in many neurodegenerative diseases (Ashleigh et al., 2023; DAlessandro et al., 2025), the identification of mitochondrial VDAC proteins as STX-interacting candidates provides a plausible molecular connection between STX signaling and mitochondrial regulation. The current results are congruent with previous findings that STX is neuroprotective against β-amyloid toxicity both in neuronal cultures and transgenic mouse models of Alzheimer’s disease, in part by supporting mitochondrial function and synaptic integrity (Gray et al., 2016; Quinn et al., 2022; Lee et al., 2025). Importantly, the current study was designed to identify molecular targets of STX and characterize the functional consequences of STX–VDAC interactions rather than to test the requirement for VDAC signaling in AD pathogenesis. Our findings therefore establish three levels of evidence: (1) the experimentally demonstrated interaction between STX and VDAC proteins; (2) the functional effects of STX on VDAC2 channel properties and mitochondrial bioenergetics; and (3) the potential, but presently unproven, relevance of this mechanism to AD-associated mitochondrial dysfunction. Whether the beneficial effects of STX previously observed in 5xFAD mice require VDAC signaling remains to be determined directly.

The loss of mitochondrial function is clearly involved in neuronal dysfunction throughout the course of AD (Ashleigh et al., 2023; DAlessandro et al., 2025). Hence, our results suggest that the neuroprotective effects of STX likely include the regulation of VDAC function in both the healthy and diseased brain. Interestingly, *Vdac2* was the most highly expressed isoform in POMC neurons, unlike the dominance of VDAC1 expression in most tissues with the exception of the testes and ovaries (Zinghirino et al., 2021). Therefore, we investigated the interaction of STX with VDAC2 using single-channel electrophysiological recordings to determine how nanomolar concentrations of STX affect the VDAC2 channel properties. In these experiments, we reconstituted VDAC2 channels in planar lipid membranes and analyzed their voltage-gating behavior, which is characterized by transitions between open and low-conducting, or closed states, in a voltage-dependent manner, a common feature of all VDAC isoforms (Queralt-Martín et al., 2020; Rosencrans et al., 2025). Nanomolar concentrations of STX increased the voltage sensitivity of VDAC2, reducing the characteristic gating voltage by about 40 mV, as observed upon the application of voltage ramps. Furthermore, in multichannel recordings, STX enhanced voltage-dependent gating in a concentration-dependent manner with a K_d_ similar to the concentrations associated with its potent effects on POMC neurons *ex vivo* (Qiu et al., 2006). Therefore, these experiments provide direct functional evidence that STX can modify VDAC2 channel behavior in a minimal membrane system, independent of other cellular proteins or transcriptional mechanisms.

These results contrast with the neurosteroid allopregnanolone, which is a potent allosteric modulator of GABA_A_ channel gating (Majewska et al., 1986; Lambert et al., 1995). Although allopregnanolone can also bind to VDACs, based on photoaffinity labeling, it does not affect their gating properties (Cheng et al., 2019). The present results indicate that STX can function as a potent enhancer of VDAC gating, independent of other protein interactions. Moreover, the STX-induced shift of ion selectivity of VDAC2 closed states towards anions further confirms a direct STX-VDAC2 interaction. The absence of measurable effects on VDAC2 open state conductance or selectivity suggests that the hydrophobic STX interacts with VDAC2 at the protein-lipid interface (Rostovtseva et al., 2020; Rovini et al., 2020).

We also investigated the effects of STX on mitochondrial physiology using mHypo43 POMC cells, which were used for the photo-affinity labeling experiments, and IHH cells, which have been established as an effective assay for mitochondrial function (Lim et al., 2008). Hepatocytes exhibit the highest mitochondrial density within the human body (Xu et al., 2023). Based on the K_d_ values obtained from our biophysical measurements of channel gating, we evaluated the effects of nanomolar concentrations of STX on mitochondrial respiration and glycolytic function in both cell lines. Notably, we observed the types of changes expected following administration of agents targeting the electron transport chain (*e.g*., oligomycin). Specifically, STX (1 nM) potently increased basal respiration, spare respiratory capacity, ATP production and proton leak in mHypo43 cells. These effects indicate enhanced mitochondrial respiratory activity and metabolic reserve. Importantly, STX also increased glycolytic function in the mHypo43 POMC neural cell line, which is congruent with the fact that neurons require glucose uptake and glycolysis to support their high energy needs (Li et al., 2023). STX was equally effective in both serum-free conditions, which eliminates endogenous estrogens, and media that included serum. These results suggest that STX is more potent than E2 in its ability to enhance mitochondrial respiratory activity and affect VDAC gating through its direct interaction, as suggested by their prominent photo-affinity labeling with BF-STX.

In general, the effects of E2 on mitochondrial function are thought to be transduced via estrogen receptors α and β (ERα, ERβ), which regulate transcriptional responses to enhance the efficiency of respiratory chain complexes and improve ATP synthesis capacity (Simpkins et al., 2008; Huang et al., 2025). Immunoreactive ERα and ERβ have been identified within mitochondria (Simpkins et al., 2008; Klinge, 2020), which is consistent with the supposition that many if not all of the actions of E2 are transcriptionally mediated. E2 activates mitochondrial biogenesis, inhibits the accumulation of reactive oxygen species (ROS), and stabilizes the mitochondrial membrane potential (Klinge, 2020). STX, in contrast, does not bind to ERα or ERβ (Qiu et al., 2003), and its effects at both the plasma membrane and mitochondrial membrane are more rapid (<15 min) than a transcriptionally mediated response (Qiu et al., 2003; Smith et al., 2013; Conde et al., 2016). Most importantly, STX directly modulates VDAC gating in the absence of any transcription factors. Notably, STX potently (1 nM) increased both basal and stress-induced (spare capacity) oxidative phosphorylation in mHypo43 and IHH cells, whereas E2 has not been found to affect basal respiration in mammalian cells (Simpkins et al., 2008). These findings identify a rapid, nonclassical pathway through which STX can regulate mitochondrial function and distinguish this mechanism from the predominantly estrogen receptor-dependent actions attributed to E2.

Several limitations of our study should be considered. First, the chemoproteomic and Seahorse experiments relied substantially on immortalized cell models. Although these approaches provided sufficient material and experimental controls, and were complemented by single-cell analysis of identified POMC neurons and electrophysiological characterization of recombinant VDAC channels, validation of the STX–VDAC mechanism in primary neurons and *in vivo* will be important. In addition, the BF-STX chemoproteomic approach identifies STX-interacting proteins but does not resolve the precise STX-binding site on VDAC. Future structural, mutagenesis, and direct binding studies will be needed to define this interaction and determine potential isoform specificity. Second, although our convergent evidence supports interaction between STX and VDAC proteins, it does not establish that VDAC2 alone is necessary for the bioenergetic or neuroprotective actions of STX. Conditional or neuron-specific loss-of-function approaches, together with evaluation of VDAC1 and VDAC3, will be needed to establish causality and address potential functional redundancy among VDAC isoforms. Likewise, the present experiments do not directly test STX–VDAC signaling in an AD model or determine whether this mechanism varies with sex or hormonal state. These questions will require targeted manipulation of VDAC isoforms in relevant neuronal populations and disease models. Finally, the non-linear concentration-response at higher STX concentrations should not be interpreted as a simple monotonic pharmacological effect. One possibility is that STX engages multiple mitochondrial targets with different apparent affinities and opposing effects on respiration. At lower concentrations, STX may preferentially engage a higher-affinity target, potentially VDAC, whereas higher concentrations may recruit lower-affinity targets that constrain the VDAC-associated increase in mitochondrial respiration. Consistent with this possibility, our BF-STX dataset identified several mitochondrial proteins involved in oxidative phosphorylation and metabolite transport, including ATP5F1C, NNT, SLC25A4, SLC25A5 and SLC25A3. ATP5F1C is required for efficient mitochondrial ATP production (Fiorillo et al., 2021), and estrogenic regulation of ATP synthase has been reported (Massart et al., 2002; Moreno et al., 2013). NNT, SLC25A4, SLC25A5 and SLC25A3 also regulate mitochondrial respiration, redox balance, and ATP production (Mayr et al., 2007; Lopert and Patel, 2014). However, BF-STX enrichment does not establish direct STX binding, relative affinity, or functional modulation of these proteins. We therefore consider this multi-target explanation a hypothesis rather than an established mechanism. Direct binding and concentration-dependent target-engagement studies will be required to determine whether the non-linear response reflects recruitment of a lower-affinity mitochondrial target, concentration-dependent effects on VDAC itself, or downstream mitochondrial feedback.

The potential relevance of these findings to neurodegenerative disease is particularly intriguing in the context of menopause, during which marked changes in brain oxidative metabolism have been proposed to contribute to AD vulnerability (Brinton et al., 2015; Mosconi et al., 2018). In neuroblastoma cell models and primary hippocampal neuronal cultures, STX protects against Aβ-associated mitochondrial dysfunction and synaptic loss (Gray et al., 2016). Moreover, Quinn and colleagues have shown that STX treatment reduces Aβ burden in 5XFAD (AD) mice treated from 6 to 8 months of life (Quinn et al., 2022). Together, these previous observations establish disease-relevant effects of STX but do not identify the molecular target responsible for those effects. The present findings provide a potential molecular framework for these previously observed actions by identifying VDAC proteins as mitochondrial STX-interacting targets and by demonstrating that STX can directly alter VDAC2 channel properties and cellular bioenergetics. This framework provides a foundation for future loss-of-function, isoform-comparison and disease-model studies to determine whether VDAC signaling mediates the neuroprotective effects of STX *in vivo*.

## MATERIAL AND METHODS

### Animals

All the animal procedures described in this study were performed in accordance with institutional guidelines based on National Institutes of Health standards and were approved by the Institutional Animal Care and Use Committee at Oregon Health and Science University (IACUC Protocol TR03-IP00000585).

*Pomc^EGFP^* (RRID: IMSR_JAX: 009593, strain of origin: C57BL/6 J) (Cowley et al., 2001) mice were used. All animals were maintained under controlled temperature and photoperiod (lights on at 0600 h and off at 1800 h) and given free access to food and water.

### mHypo43 (mHypoE-N43/5) cells

The embryonic mouse hypothalamus cell line N43/5 (mHypoE-N43/5) was obtained from Cellutions Biosystems (Cedarlane, cat. no. CLU127) and cultured according to the supplier’s protocol. Cells were grown in 10 cm tissue culture dishes at 37 °C in 5% CO₂ using DMEM (Gibco, cat. no. 11960-044) containing 4.5 mg/mL D-glucose, supplemented with 2 mM L-glutamine (Sigma-Aldrich, cat. no. G7513), 1× non-essential amino acids (Gibco, cat. no. 11140-050), 1% penicillin/streptomycin (Sigma-Aldrich, cat. no. P4333), and 10% FBS. Freshly thawed cells were initially seeded and allowed to attach for 4–6 h before medium replacement. Cells were passaged at 70–90% confluence using trypsin and split at ratios between 1:5 and 1:10.

### Synthesis and physiological validation of BF-STX probe

The novel bifunctional STX derivative (BF-STX) containing both a photo-cross-linkable diazirine group and an alkyne handle was designed by the Schultz Lab and synthesized in collaboration with SiChem GmbH (Germany) (**Figure 1A** and see Supplementary Information). To demonstrate that this construct was functional, we tested the efficacy of BF-STX to modulate the excitability of POMC neurons in *ex vivo* electrophysiological slice experiments. Indeed, BF-STX was as efficacious as STX in increasing POMC cell excitability (Qiu et al., 2003; Qiu et al., 2006) (**Figure 1B**).

#### Confocal imaging of BF-STX labelling in brain slices

Coronal hypothalamic arcuate slices (250 µm thick) containing POMC neurons were incubated with 10 µM BF-STX for 30 min, then subjected to UV photo-crosslinking (2.5 min, 1000 W Xe lamp with a 350 nm high-pass filter). Slices were fixed in cold methanol (–20 °C, 20 min), and BF-STX–protein conjugates were visualized by Cu(I)-catalyzed click chemistry using Alexa Fluor 488 picolyl azide. Nuclei were counterstained with DAPI. Imaging was performed on an Olympus FV1200 confocal microscope with a 60× oil-immersion objective, using the third ventricle (3V) as an anatomical landmark (**Figure 1C**).

#### Confocal Imaging of BF-STX labelling in mHypo43 cells

mHypo43 (POMC neurons) were treated with 10 µM BF-STX for 30 min, followed by UV crosslinking under the same conditions (2.5 min, 1000 W Xe lamp, 350 nm filter) (**Figure 1D**). After fixation in cold methanol (–20 °C, 20 min), cells were labeled with Alexa Fluor 488 picolyl azide and counterstained with DAPI. Confocal imaging (Olympus FV1200, 60× oil objective) showed robust intracellular accumulation of BF-STX in mHypo43.

### Visualized whole-cell patch recording

Whole-cell current clamp and voltage clamp recordings were made from POMC neurons as previously described (Qiu et al., 2010; Qiu et al., 2014). Coronal arcuate slices (250 μm) were prepared from gonadectomized females, 10 weeks or older as previously described (Qiu et al., 2010; Qiu et al., 2014). The slices were then transferred to an auxiliary chamber in which they were kept at room temperature (25 °C) in artificial CSF (aCSF) consisting of the following (in mm): 124 NaCl, 5 KCl, 2.6 NaH_2_PO_4_, 2 MgSO_4_, 2 CaCl_2_, 26 NaHCO_3_, 10 HEPES, 10 glucose, pH 7.4, until recording (recovery for 2 h). A single slice was transferred to the recording chamber at a time and was kept viable by continually perfusing with warm (35 °C), oxygenated aCSF at 1.25 ml/min. Whole-cell patch recordings were made from POMC neurons using an Olympus BX51 W1 fixed -stage microscope equipped with epifluorescence and infrared-differential interference contrast (IR-DIC) video microscopy. Patch pipettes (A-M Systems; 1.5 μm outer diameter borosilicate glass) were pulled on a Brown/Flaming puller (Sutter Instrument, model P-97) and filled with the following solution: 128 mM potassium gluconate, 10 mM NaCl, 1 mM MgCl_2_, 11 mM EGTA, 10 mM HEPES, 3 mM ATP, and 0.25 mM GTP adjusted to pH 7.3 with KOH; 295 mOsm. Pipette resistances ranged from 3.5 to 4 MΩ. In whole-cell configuration, access resistance was less than 30 MΩ; the access resistance was 80% compensated. The input resistance was calculated by measuring the slope of the I–V relationship curve between −70 and −50 mV. Standard whole-cell patch recording procedures and pharmacological testing were performed as previously described (Qiu et al., 2010). Electrophysiological signals were digitized with a Digidata 1322 A (Axon Instruments) and the data were analyzed using p-Clamp software (Molecular Devices, Foster City, CA).

To study the effect of BF-STX on µ-opioid receptor-mediated responses in POMC neurons, a drug administration protocol was employed during whole-cell patch-clamp recordings in voltage-clamp mode (V_hold_ = –60 mV). After forming a gigaseal and achieving the whole-cell configuration, slices were perfused with tetrodotoxin (TTX, 1 μM) for 5 minutes in order to synaptically isolate the neurons. The µ-opioid receptor-mediated response was evoked by perfusing [D-Ala2, N-MePhe4, Gly-ol]-enkephalin (DAMGO, 300 nM) until a steady-state outward current was reached. Following drug washout, the current returned to its pre-drug baseline. Cells were then treated with BF-STX for 15 minutes. DAMGO was perfused again, and the second response was recorded.

A standard artificial cerebrospinal fluid was used (Qiu et al., 2010). TTX (1 mM, Alomone Labs) and DAMGO (1 mM, d-Ala2, N-Me-Phe4, Gly-ol5-enkephalin; Peninsula Laboratories, Inc., Belmont, CA) were dissolved in H_2_O. BF-STX (10 mM) was dissolved in 100% ethanol. Aliquots of the stock solutions were stored as appropriate until needed.

### Identification of candidate STX targets in POMC neurons using BF-STX

mHypo43 cells (3 x 10^6^ cells) were grown in culture medium (see above) and incubated with 10 μM BF-STX in DMEM for 5 min or 30 min. Photo-crosslinking was performed for 5 min. BF-STX without UV crosslinking and pre-incubation with STX before cross-linking served as negative controls. After washing and scraping, cells were lysed on ice by probe sonication (three 15 s bursts). Lysates were subjected to copper-catalyzed azide–alkyne cycloaddition (CuAAC) with picolyl azide agarose beads for selective enrichment of labeled proteins. Azide beads (200 µL) were washed in deionized water, added to lysate with CuSO₄ (1 mM), sodium ascorbate (1 mM), and TBTA (100 µM), and incubated at room temperature for 1 h with rotation. Beads were pelleted (1000 x g, 2 min), transferred to centrifuge columns, and washed with PBS (3×), bead wash buffer 1 (100 mM Tris-HCl, pH 8.0, 250 mM NaCl, 5 mM EDTA, 1% SDS; 5×), and bead wash buffer 2 (100 mM Tris-HCl, pH 8.0, 8 M urea; 10×). Beads were transferred to clean tubes and pelleted.

Captured proteins were reduced in digestion buffer (100 mM Tris-HCl, pH 8.0, 2 mM CaCl₂, 10% acetonitrile) with DTT (10 mM) at 42 °C for 30 min and alkylated with iodoacetamide (40 mM) at room temperature in the dark for 30 min. Bead-bound proteins were digested overnight at 37 °C with sequencing-grade trypsin. Peptides were desalted on C18 cartridges and stored at −80 °C prior to shipping to EMBL.

#### Identification of isolated proteins by LC-MS/MS (see Supplementary Information for complete protocol)

Up to 10 µg of peptides were labeled using TMTpro™ 16plex reagent as previously described (Thompson et al., 2019). Briefly, 0.5 mg of TMT reagent was dissolved in 45 µL of 100% acetonitrile. Subsequently, 4 µL of this solution was added to each peptide sample, followed by incubation at room temperature for 1 hour. The labeling reaction was quenched by adding 4 µL of a 5% aqueous hydroxylamine solution and incubating for an additional 15 minutes at room temperature. Labeled samples were then combined for multiplexing, desalted using an Oasis® HLB µElution Plate (Waters) according to the manufacturer’s instructions, and dried by vacuum centrifugation.

Peptides were separated on an UltiMate 3000 RSLCnano system using a C18 trapping cartridge (µ-Precolumn PepMap™ 100, 300 µm × 5 mm, 5 µm, 100 Å) and an analytical column (nanoEase™ M/Z HSS T3, 75 µm × 250 mm, 1.8 µm, 100 Å). Samples were trapped at 30 µL/min in 0.05% TFA for 6 min and then gradient-eluted at 0.3 µL/min with solvent A (3% DMSO, 0.1% formic acid in water) and solvent B (3% DMSO, 0.1% formic acid in acetonitrile): 2–8% B (4 min), 8–26% B (104 min), 26–38% B (4 min), 40–80% B (0.1 min), 80% B (3.9 min), followed by re-equilibration to 2% B (4 min).

Peptides were analyzed on an Orbitrap Fusion™ Lumos™ Tribrid™ mass spectrometer in positive ion mode with a Pico-Tip nanospray emitter (2.2 kV spray voltage, capillary temperature 275 °C). Full MS scans (m/z 375–1500) were acquired in the Orbitrap at 120 000 resolution with a maximum injection time of 50 ms. Data-dependent MS/MS spectra were acquired at 30 000 resolution with a maximum injection time of 94 ms and AGC target of 200%. HCD fragmentation was used with normalized collision energy of 34%, a quadrupole isolation window of 0.7 m/z, and dynamic exclusion of 60 s; precursor charge states 2–7 were selected for fragmentation.

#### Data analysis

Raw files were converted to mzML format using MSConvert (ProteoWizard, RRID:SCR_012056) with peak picking and zlib compression. Spectra were searched with MSFragger in FragPipe (v19.0) against the UniProt (RRID:SCR_002380) Homo sapiens reference proteome (UP000005640) including common contaminants and reversed sequences. Carbamidomethylation (C, 57.0215) and TMTpro (K, 304.2072) were set as fixed modifications; oxidation (M, 15.9949), protein N-terminal acetylation (42.0106), and TMTpro (peptide N-terminus, 304.2072) were included as variable modifications. Trypsin specificity with up to two missed cleavages and a minimum peptide length of seven residues was used; precursor and fragment mass tolerances were set to 20 ppm; peptide and protein FDR were controlled at 1% using standard FragPipe settings.

FragPipe (protein.tsv files) were processed using the R programming environment (ISBN 3-900051-07-0). Initial data processing included filtering out contaminants and reverse proteins. Only proteins quantified with at least 2 unique peptides (Unique.Peptides >= 2) were considered for further analysis. 878 proteins passed the quality control filters. In order to correct for technical variability, batch effects were removed using the ’removeBatchEffect’ function of the limma (RRID:SCR_010943) package (Ritchie et al., 2015) on the log2 transformed raw TMT reporter ion intensities (’channel’ columns). Subsequently, normalization was performed using the ’normalizeVSN’ function of the limma package (VSN - variance stabilization normalization – (Huber et al., 2002)). For visualization, proteins abundances were normalized to the control condition by dividing by the median of control replicates (’ctrl’ column). These control ratios were used for clustering and heatmap display. Differential expression analysis was performed using the moderated t-test provided by the limma package (Ritchie et al., 2015). The model accounted for replicate information by including it as a factor in the design matrix passed to the ’lmFit’ function. To obtain p-values and false discovery rates (FDRs), the ’fdrtool’ function from the fdrtool package (Strimmer, 2008) was used to analyze the t-values produced by limma for certain comparisons. Proteins were annotated as hits if they had a false discovery rate (FDR) below 0.05 and an absolute fold change greater than 2. Proteins were considered candidates if they had an FDR below 0.2 and an absolute fold change greater than 1.5. Clustering based on the median protein abundances normalized by median of control condition was conducted to identify groups of proteins with similar patterns across conditions. The ’kmeans’ method was employed, using Euclidean distance as the distance metric and ’ward.D2’ linkage for hierarchical clustering. The optimal number of clusters (6) was determined using a hybrid Silhouette approach. The threshold was set as the maximum of either the 80th percentile of silhouette values or a fixed floor of 0.25 (threshold = 0.403), selecting the maximum number of clusters exceeding this threshold. This hybrid approach balances data-driven cluster selection while avoiding selection from noisy low-quality regions.

### Cell harvesting of dispersed and POMC neurons and quantitative real-time PCR (qPCR)

Cell harvesting and qPCR was conducted as previously described (Bosch et al., 2013). The ARH was microdissected from basal hypothalamic coronal slices obtained from female *Pomc^EGFP^* mice in which POMC neurons were labeled with YFP (n = 3 animals/group). The dispersed cells were visualized, patched, and then harvested (10 cells/tube) as described previously (Bosch et al., 2013). Briefly, the tissue was incubated in papain (7 mg/ml in oxygenated aCSF) for 50 min at 37°C and washed 4 times in low Ca^2+^ aCSF and two times in aCSF. Gentle trituration with Pasteur pipettes was used to disperse the neurons onto a glass-bottom dish. Oxygenated aCSF circulated into the plate keeping the cells clear of debris. Only healthy cells with processes and a smooth cell membrane were harvested. The cells were harvested using the XenoWorks Microinjector System (Sutter Instruments, Navato, CA), which provided negative pressure in the pipette and fine control to draw the cell up into the pipette. Cells were harvested as pools of 10 individual cells/tube.

Primers for the genes that encode for *Vdac1, Vdac2, Vdac3* and *β-actin* were designed using Clone Manager software (Sci Ed Software) (RRID:SCR_014521) to cross at least one intron-exon boundary and optimized as previously described using Power SYBR Green method (Bosch et al., 2013; Nestor et al., 2016). We have published previously the primers for *β-actin* (Qiu et al., 2018), which have similar efficiency as the *Vdac* primers (see Table1). Controls included neuronal pools reacted without reverse transcriptase (RT), hypothalamic RNA reacted with RT and without RT, as well as water blanks. Primers (**Table 1**) for qPCR were further tested for efficiency (E = 10^(−1/m)^ – 1) (Livak and Schmittgen, 2001; Pfaffl, 2001; Bosch et al., 2013).

**Table 1.** Primer Table.

| Gene Name<br>(encodes for) | Accession<br>Number | Primer<br>Location (nt) | Product<br>Length (bp) | Annealing<br>Temp (°C) | <u>Efficiency</u> |  |  |
| --- | --- | --- | --- | --- | --- | --- | --- |
|  |  |  |  |  | Slope | r<sup>2</sup> | % |
| VDAC1 | NM_001362693 | 233-352 | 120 | 60 | -3.445 | 0.86 | 95 |
| VDAC2 | NM_011695 | 192-266 | 77 | 60 | -3.567 | 0.99 | 93 |
| VDAC3 | NM_00119898 | 905-987 | 83 | 60 | -3.507 | 0.99 | 91 |
| β-Actin | NM_007393 | 446-555 | 110 | 60 | -3.465 | 0.99 | 95 |

qPCR was performed on a Quantstudio 7 Flex Real-Time PCR System (Life Technologies) using Power SYBR Green (Life Technologies) Master Mix according to established protocols (Bosch et al., 2013). The comparative ΔΔC_T_ method (Livak and Schmittgen, 2001; Pfaffl, 2001) was used to determine values from duplicate samples of 4 μl for the target genes and 2 μl for the reference gene β-actin in a 20 μl reaction volume containing 1× Power SYBR Green PCR Master Mix (Applied Biosystems) and 0.5 μM forward and reverse primers. Three 10 cell neuronal pools/animal (n=3 animals) were used to determine the relative linear quantity using the 2^-ΔΔCT^ equation (Bosch et al., 2013). In order to compare the relative expression levels of target genes *Vdac1*, *Vdac2* and *Vdac3* in POMC^EGFP^ neurons the mean Δ CT for target gene *Vdac2* was used as the calibrator.

### Measurements of VDAC activity in planar lipid membranes

Planar lipid bilayers were made by the apposition of two monolayers comprised of a 1:1 (w:w) mixture of dioleoyl-phosphatidylcholine (DOPC) and dioleoyl-phosphatidylethanolamine (DOPE) lipids (1:1 w/w) or diphytanoyl-phosphatidylcholine (DPhPC) (Avanti Polar Lipids, Alabaster, AL) dissolved in pentane. These bilayers were formed across a ∼125 μm (in multichannel experiments) and 75 μm (in single-channel experiments) diameter aperture in the 15-μm-thick Teflon partition separating two ∼1.5-mL compartments, as previously described (Rostovtseva et al., 2006). The recombinant human VDAC2 was a generous gift of Tsyr-Yan Dharma Yu (Institute of Atomic and Molecular Sciences, Academia Sinica, Taipei, Taiwan). VDAC2 channel insertion was achieved by the addition of 0.2-0.5 µl of VDAC2 solution diluted in 100 mM Tris, 50 mM KCl, 1 mM EDTA, 15% (v/v) DMSO, 2.5% (v/v) Triton X-100, pH 7.35 to the cis (grounded) compartment of the experimental chamber while stirring, as described previously (Rostovtseva et al., 2006; Rostovtseva and Bezrukov, 2015).). Ion current recordings were acquired using an Axopatch 200B amplifier (Axon Instruments) in the voltage-clamp mode, following previously reported protocols (Rostovtseva and Bezrukov, 2015). Current signals were filtered at 10 kHz with a low-pass Bessel (8 pole) filter, digitized at a sampling frequency of 50 kHz, and analyzed using pClamp10.7 software (Axon Instruments For single-channel analyses in Clampfit 10.7, recordings were further digitally filtered with a 5 kHz 8-pole Bessel low-pass filter. Potential is defined as positive when it is greater on the cis side, the side of compound STX and VDAC2 addition. STX was added from its stock solution in DMSO.

Voltage dependence of VDAC2 gating was analyzed using a previously described protocol (Rostovtseva et al., 2006; Teijido et al., 2014) under the application of a slow, symmetric triangular voltage wave (±60 mV, 5 mHz) generated by an Arbitrary Waveform Generator 33220A (Agilent Technologies). Data were sampled at 2 Hz frequency and analyzed as detailed earlier (Teijido et al., 2014) using a custom in-house algorithm (Rappaport et al., 2015) and pClamp 10.7 software (RRID:SCR_011323). In each experiment, current records were collected from membranes containing more than 20 channels in response to 5–10 periods of voltage waves to ensure data collection from more than 100 channels per experiment. Only the part of the wave, during which the channels were reopening, was used for the subsequent analysis for the reasons described elsewhere (Colombini, 1989; Teijido et al., 2014). G_min_, the minimal multichannel conductance in multichannel experiments, was measured at applied potentials in the range of 50 mV < |V| < 60 mV.

VDAC selectivity was measured in 1M (cis) / 200mM (trans) KCl gradient buffered with 5 mM HEPES at pH 7.4, as described previously (Teijido et al., 2014). In all experiments, VDAC2 was added to the cis compartment. VDAC2 ion selectivity was calculated from the potential corresponding to the zero current (Ψ_rev_) using a linear regression fit to the data.

### Flow cytometry assay

mHypo43 cells were seeded in 24-well plates at 1x10^5^ cells/well and incubated overnight in serum-free DMEM without phenol red in the presence of 50 nM or 100 nM STX. 1 µM LIVE/DEAD^TM^ Fixable Near-IR Dead Cell stain (Thermo Fisher Scientific, L34976) was added to each sample and the fluorescence was measured by exciting cells with 637 nm laser and detecting emission with 780/60 nm bandpass filter. Labeled cells were acquired on a flow cytometer (BD FACSymphony A5) with FACSDiva software v9.3.1(RRID:SCR_001456). Analysis was performed in FlowJo v10.10 (RRID:SCR_008520) with gating on singlet live cells, and the geometric mean for label fluorescence was determined for each condition.

#### Mitochondrial membrane potential

Cells were trypsinized and incubated with 5nM TMRM (Tetramethylrhodamine methyl ester, Thermo Fisher Scientific, M20036) and LIVE/DEAD^TM^ Fixable Near-IR Dead Cell stain in PBS for 30 mins at 37 °C in a 5% CO_2_ incubator. Cells were washed and resuspended in 250 µL PBS and acquired on low speed. The fluorescence of TMRM was measured by exciting the cells with a 561 nm laser, and the emission was detected with bandpass filter of 586/15 nm.

#### Mitochondrial calcium

Cells were trypsinized and incubated with 2 µM Rhod2-AM (Thermo Fisher Scientific, R1245MP) and 0.02% Pluronic^®^ F-127 (Sigma-Aldrich, P2443) in PBS for 45 mins at room temperature. Cells were washed and incubated with LIVE/DEAD^TM^ Fixable Near-IR Dead Cell stain in 250 µL PBS for 30 mins at room temperature and acquired on low speed. The fluorescence of Rhod2 was measured by exciting the cells with a 561 nm laser, and the emission was detected with bandpass filter of 586/15 nm.

#### Mitochondrial ATP

Cells were trypsinized and incubated with 5 µM Biotracker ATP-Red (Sigma-Aldrich, SCT045) and LIVE/DEAD^TM^ Fixable Near-IR Dead Cell stain in PBS for 15 mins at 37 °C and 5% CO_2_. Cells were washed and resuspended in 250 µL PBS and acquired on low speed. The fluorescence of ATP-Red was measured by exciting cells with a 561 nm laser, and the emission was detected with a bandpass filter of 586/15 nm.

### Seahorse Assays

Cellular bioenergetics were assessed using the Seahorse XFe96 Analyzer (Agilent Technologies) with either the XF Cell Mito Stress Test Kit (Agilent Technologies, 103015-100) or the XF Glycolysis Stress Test Kit (Agilent Technologies, 103020-100), following the manufacturer’s protocols. Seahorse XFe 96-well plates (Agilent Technologies, 103794-100) were coated with rat tail collagen type I (Gibco, A10483-01) for 1 h at room temperature, air-dried under sterile conditions, and seeded with immortalized human hepatocytes (IHH) or mHypo43 cells at 20,000 cells per well, followed by overnight incubation at 37°C with 5% CO₂. The Seahorse XFe96 sensor cartridge (Agilent Technologies, W11425) was hydrated overnight in Seahorse XF Calibrant (Agilent Technologies, 100840-000) at 37°C in a non-CO₂ incubator. On the day of each assay, cells were washed twice with Dulbecco’s phosphate-buffered saline (Gibco, 14190-144), serum-starved for 6 h, and then washed twice with Seahorse XF DMEM medium, pH 7.4 (Agilent Technologies, 103575-100). Cells were incubated for 1 h at 37°C in a non-CO₂ incubator with either 1 nM STX, 5nM STX (for mHypo43 cells), 10 nM STX, or Seahorse XF assay medium alone (control) immediately prior to the assay. To assess the impact of serum starvation, additional assays were performed in the same manner but without the 6 h serum starvation step.

For the Mito Stress Test, the Seahorse assay medium was supplemented with 10 mM glucose, 1 mM pyruvate, and 2 mM L-glutamine, and injection ports were loaded with 1.5 µM oligomycin, 1 µM FCCP, and 0.5 µM rotenone/antimycin A. Oxygen consumption rate (OCR) was measured in real time using cycles of mixing (3 min), waiting (2 min), and measuring (3 min), and basal respiration, ATP-linked respiration, maximal respiration, spare respiratory capacity, and non-mitochondrial respiration were calculated.

For the Glycolysis Stress Test, the Seahorse assay medium was supplemented with 2 mM L-glutamine only, and injection ports were loaded with 10 mM glucose, 1 µM oligomycin, and 50 mM 2-deoxy-D-glucose (2-DG). Extracellular acidification rate (ECAR) was measured using the same cycle parameters described above, and glycolysis, glycolytic capacity, glycolytic reserve, and non-glycolytic acidification were calculated.

All data were analyzed using Wave software (Agilent Technologies). Each treatment group was assayed in at least eight technical replicates per plate, and all experiments were independently repeated a minimum of three times. Wells with poor adherence or OCR/ECAR values exceeding two standard deviations from the mean were excluded from analysis.

Data are presented as mean ± standard error (SE). Statistical significance was determined using one-way ANOVA followed by Dunnett’s multiple comparison test.

## Additional Files

Additional experimental details, materials, methods and supporting figures, including NMR spectra, are available in the Supplemental Information pdf file.

## Note

The authors declare no competing financial interests.

## ACKNOWLEDGEMENTS

The authors would like to thank Jin Hui-Deng for his assistance in breeding and genotyping the transgenic mouse lines at OHSU. The click chemistry, molecular biological and electrophysiological studies were supported by the National Institutes of Health grants: NIA grant R21 AG080057 (MJK & CS) and NIDDK grant R01 DK068098 (MJK & OKR). Research at the Intramural Research Program of the *Eunice Kennedy Shriver* National Institute of Child Health and Human Development, National Institutes of Health was funded by NIH grant ZIA HD000072-18 (SMB). The contributions of the NIH authors (SMB, TKR, MR, MW, WF) are considered Works of the United States Government. The findings and conclusions presented in this paper are those of the authors and do not necessarily reflect the views of the National Institutes of Health or the U.S. Department of Health and Human Services.

## I) General Procedures

All reactions were carried out with dry solvents under an argon atmosphere, unless otherwise noted. Anhydrous methylene chloride (CH_2_Cl_2_), anhydrous N,N-dimethylformamide (DMF), N,N-diisopropylethylamine (iPr_2_NEt), triethylamine (Et_3_N), triphenylphosphine (Ph_3_P), trichloroacetonitrile (CCl_3_CN) and HCl 4M in dioxane were purchased from Merck and used as received. Building block **S1** was purchased from Fluorochem and used as received. Building Block **S2** was purchased from Enamine and used as received. Building block **S5** was synthesized according to a previously reported procedure (Thomas S. Scanlan, Toru Iijima. Selective estrogen receptor modulator compositions and methods for treatment of disease. WO 2009062082 A2, 14 May **2009**. US8236987B2). Ethyl acetate (EtOAc), TFA (CF_3_CO2H), methanol (MeOH) and water (H_2_O) were purchased at the highest commercial quality and used without further purification. Yields refer to chromatographically homogeneous materials, unless otherwise stated. Reactions that required heating were operated on a magnetic stirrer with an oil bath. Reactions and purity of final product BF-STX were monitored via high performance liquid chromatography (HPLC) analyses, performed on a Merck-Hitachi L-6000 pump, interfaced with a Merck-Hitachi L-4000A UV detector at 254 nm. **Method A1**: Column (YMC Triart C18-S RP, 120A, 10 µm, 250 mm x 4 mm), Eluent (65% MeOH + 35% water + 0.1% TFA), Flow (1.5 ml/min), Isocratic. **Method A2**: Column (YMC Triart *C18-S RP, 120A, 10 µm, 250 mm x 4 mm), Eluent (80% MeOH + 20% water + 50 mM TEAF pH 7), Flow (1.5 ml/min), Isocratic. **Method A3**: Column (YMC Triart C18-S RP, 120A, 10 µm, 250 mm x 4 mm), Eluent (75% MeOH + 25% water + 50 mM TEAF pH 7), Flow (1.5 ml/min), Isocratic. Purification of intermediates and final product were performed via preparative RP-HPLC on a Knauer K-1001 pump, interfaced with a Büchi C-630 UV detector at 254 nm. **Method P1**: Column (YMC Triart C18-S RP, 120A, 10 µm, 250 mm x 50 mm), Eluent (65% MeOH + 35% water + 0.1% TFA), Flow (50 ml/min), Isocratic. **Method P2**: Column (YMC Triart C18-S RP, 120A, 10 µm, 250 mm x 50 mm), Eluent (85% MeOH + 15% water + 50 mM TEAB pH 7), Flow (50 ml/min), Isocratic. **Method P3**: Column (YMC Triart C18-S RP, 120A, 10 µm, 250 mm x 50 mm), Eluent (75% MeOH + 25% water + 50 mM TEAB pH 7), Flow (25 ml/min), Isocratic. NMR spectra were recorded on Bruker AV-600 spectrometer. Chemical shifts (δ) are reported in parts per million (ppm) and calibrated using residual undeuterated solvent (CDCl_3_ δ = 7.27 ppm for ^1^H NMR, δ = 77.16 ppm for ^13^C NMR) as an internal reference. The following abbreviations were used to explain the multiplicities: s = singlet, d = doublet, t = triplet, q = quartet, br = broad, ddd = doublet of doublet of doublets, m = multiplet. Electrospray Ionization Mass spectra (ESI-MS) were recorded on Bruker amaZon SL ESI-ion trap mass spectrometer*.

## II) Experimental Procedures and Characterization Data of Compounds

Preparation of the Boc-protected diazirine-bearing building block **S3**:

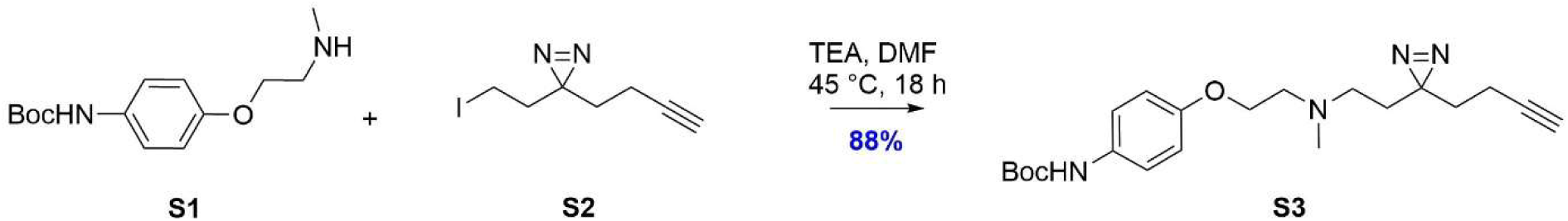

*Commercially available **S1** (0.43 g, 1.61 mmol, 1 equiv) and commercially available **S2** (0.4 g, 1.61 mmol, 1 equiv) were dissolved in 10 ml of DMF followed by the addition of TEA (335 µl, 2.42 mmol, 1.5 equiv). The solution was stirred at 45 °C for 18 h after which it was diluted with water (50 ml) and extracted with EtOAc (3 x 20 ml). The organic layers were pooled together, dried over MgSO_4_, filtered and concentrated under reduced pressure. The obtained yellow crude oil (0.77 g) was purified via RP-HPLC (method P1, retention time 16.62 min). The product-containing fraction was dried under vacuum to constant weight, affording **S3** (0.55 g, 1.42 mmol) in 88 % yield as a pale-yellow oil.* ***S3****: **^1^H NMR** (600 MHz, CDCl_3_) δ = 7.46 (d, J = 9.0 Hz, 2H), 7.03 (d, J = 8.6 Hz, 2H), 6.53 (br, 1H), 4.05 (t, J = 5.6 Hz, 2H), 2.81 (t, J = 6.0 Hz, 2H), 2.39 (t, J = 5.8 Hz, 2H), 2.34 (s, 3H), 2.00 – 1.96 (m, 3H), 1.66 – 1.63 (m, 4H), 1.42 (s, 9H); **^13^C NMR** (150 MHz, CDCl_3_) δ = 157.23 (1C), 152.35 (1C), 132.26 (1C), 123.67 (2C), 118.40 (2C), 85.25 (1C), 82.78 (1C), 66.10 (1C), 62.18 (1C), 55.98 (1C), 52.17 (1C), 42.44 (1C), 42.25 (1C), 32.40 (1C), 29.64 (1C), 22.83 (3C), 19.57 (1C); **ESI-MS** (positive mode) m/z: 387.37 [M + H]⁺*.

Preparation of the diazirine-bearing building block **S4**:

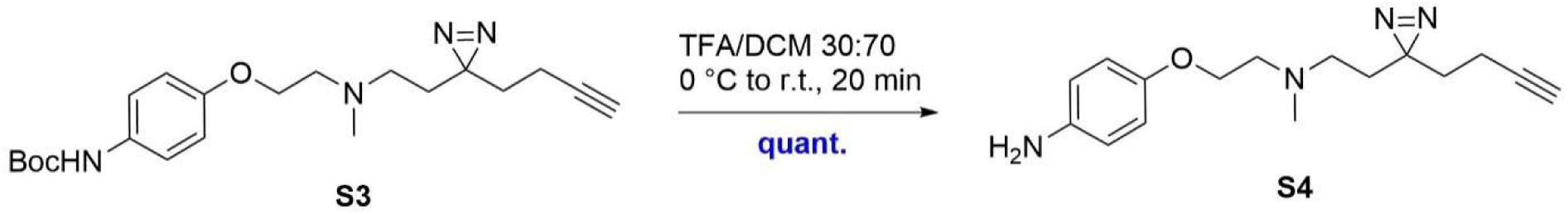

***S3*** *(0.50 g, 1.3 mmol, 1 equiv) was dissolved in DCM (14 ml) and the resulting solution was cooled down to 0 °C. TFA (6 ml) was added to the above mixture. The reaction was allowed to warm to room temperature and was then stirred for additional 20 minutes. RP-HPLC analysis (method A1, retention time 1.5 min) showed complete conversion to the desired product. The mixture was concentrated under reduced pressure affording **S4** (0.39 g, 1.3 mmol) in quantitative yield as a yellowish wax used as such in the next step.* ***S4****: **^1^H NMR** (600 MHz, CDCl_3_) δ = 7.37 (d, J = 9.0 Hz, 2H), 6.96 (d, J = 8.6 Hz, 2H), 3.97 (t, J = 5.6 Hz, 2H), 2.73 (t, J = 6.0 Hz, 2H), 2.41 (t, J = 5.8 Hz, 2H), 2.28 (s, 3H), 1.90 – 1.86 (m, 3H), 1.60 – 1.57 (m, 4H); **^13^C NMR** (150 MHz, CDCl_3_) δ = 137.85 (1C), 132.26 (1C), 122.01 (2C), 114.59 (2C), 82.78 (1C), 66.20 (1C), 66.11 (1C), 55.97 (1C), 52.15 (1C), 42.41 (1C), 42.23 (1C), 32.38 (1C), 29.63 (1C), 19.55 (1C); **ESI-MS** (positive mode) m/z: 286.27 [M + H]⁺*.

Preparation of the MOM-protected BF-STX **S6**:

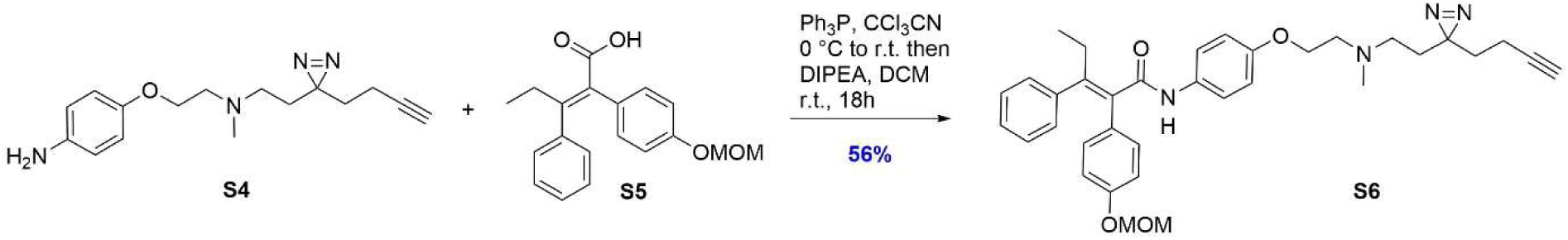

*Tetra-substituted olefin **S5** was prepared according to a previously reported procedure (Thomas S. Scanlan, Toru Iijima. Selective estrogen receptor modulator compositions and methods for treatment of disease. WO 2009062082 A2, 14 May **2009**. US8236987B2). To a solution of **S5** (0.232 g, 0.74 mmol, 1 equiv) and triphenylphosphine (0.39 g, 1.48 mmol, 2 equiv) in DCM (8 mL) at 0 °C was added trichloroacetonitrile (150 µL, 1.48 mmol, 2 equiv) dropwise. The reaction mixture was allowed to warm to room temperature and stirred for 4 h. A solution of **S4** (0.234 g, 0.82 mmol, 1.1 equiv) and DIPEA (386 µL, 2.22 mmol, 3 equiv) in DCM (4 mL) was added dropwise to the activated carboxylic acid. The mixture was stirred at room temperature for 18 h, concentrated under reduced pressure, and purified by RP-HPLC (method P2, retention time 29.40 min). The product-containing fraction was concentrated to give **S6** (0.24 g, 0.41 mmol, 56%) as a yellowish wax.* ***S6****: **^1^H NMR** (600 MHz, CDCl_3_) δ = 7.43 (d, J = 9.0 Hz, 2H), 7.22 – 7.14 (ddd, J = 13.2, 7.8, 6.1 Hz, 3H), 7.07 – 6.87 (m, 4H), 6.82 (d, J = 9.0 Hz, 2H), 6.52 (d, J = 8.6 Hz, 2H), 5.10 (s, 2H), 4.02 (t, J = 5.6 Hz, 2H), 3.44 (s, 3H), 2.79 – 2.63 (m, 4H), 2.29 – 2.17 (m, 5H), 2.02 – 1.92 (m, 3H), 1.73 – 1.67 (m, 4H), 1.05 (t, J = 7.4 Hz, 3H); **^13^C NMR** (150 MHz, CDCl_3_) δ = 171.90 (1C), 157.54 (1C), 157.08 (1C), 143.40 (1C), 141.46 (1C), 136.04 (1C), 132.93 (1C), 131.75 (2C), 130.41 (2C), 129.23 (2C), 129.10 (2C), 127.90 (1C), 123.34 (2C), 115.66 (3C), 96.02 (2C), 82.90 (1C), 69.25 (1C), 66.15 (1C), 58.15 (2C), 56.00 (1C), 52.18 (1C), 42.46 (1C), 42.27 (1C), 32.42 (1C), 30.54 (1C), 29.68 (1C), 19.58 (1C), 13.21 (1C); **ESI-MS** (positive mode) m/z: 581.40 [M + H]⁺*.

Preparation of the **BF-STX**:

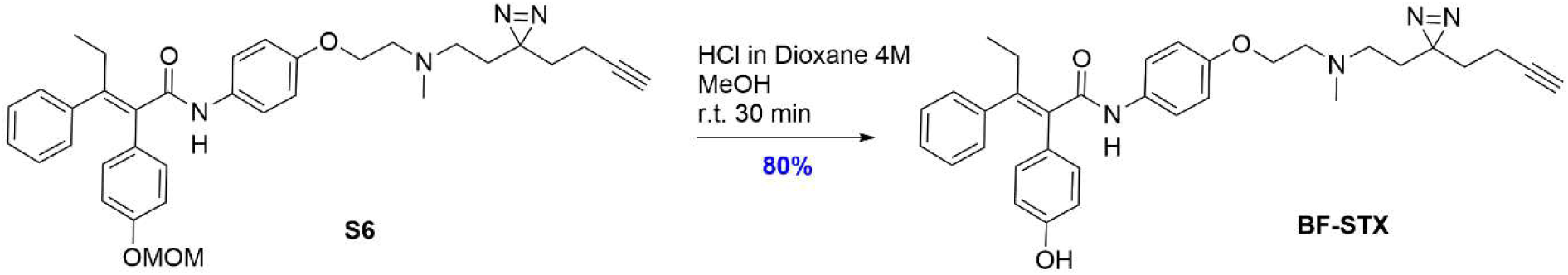

***S6*** *(0.19 g, 0.33 mmol, 1 equiv) was dissolved in MeOH (5 mL) at r.t. after which HCl 4M dissolved in dioxane (1.02 ml) was added. The resulting solution was allowed to stir at r.t. for 30 minutes. RP-HPLC analysis (method A2) showed complete conversion to the desired product. The solution was concentrated and purified via RP-HPLC (method P3) to afford BF-STX (0.141 g, 0.26 mmol) in 80 % yield as a white solid with a purity of 99.4 % as measured via RP-HPLC (method A3). **BF-STX**: **^1^H NMR** (600 MHz, CDCl_3_) δ = 7.38 (d, J = 9.0 Hz, 2H), 7.15 (ddd, J = 13.2, 7.8, 6.1 Hz, 3H), 7.09 – 6.98 (m, 2H), 6.87 (d, J = 8.6 Hz, 2H), 6.80 (d, J = 9.0 Hz, 2H), 6.54 (d, J = 8.6 Hz, 2H), 4.03 (t, J = 5.6 Hz, 2H), 2.85 – 2.68 (m, 4H), 2.43 – 2.34 (m, 2H), 2.31 (s, 3H), 2.06 – 1.90 (m, 3H), 1.64 (t, J = 7.5 Hz, 4H), 1.02 (t, J = 7.4 Hz, 3H); **^13^C NMR** (150 MHz, CDCl_3_) δ = 168.36 (1C), 156.13 (1C), 155.41 (1C), 137.85 (1C), 135.52 (1C), 132.26 (1C), 131.10 (1C), 130.44 (2C), 129.03 (2C), 129.01 (2C), 128.51 (2C), 127.62 (1C), 122.01 (2C), 114.59 (2C), 82.78 (1C), 69.23 (1C), 66.13 (1C), 55.98 (1C), 52.16 (1C), 42.45 (1C), 42.26 (1C), 32.40 (1C), 30.53 (1C), 29.65 (1C), 19.56 (1C), 13.23 (1C); **ESI-MS** (positive mode) m/z: 537.34 [M + H]⁺; m/z: 559.32 [M + Na]⁺*

## III) Supplementary Figures (S1-S6)

**Figure S1.**
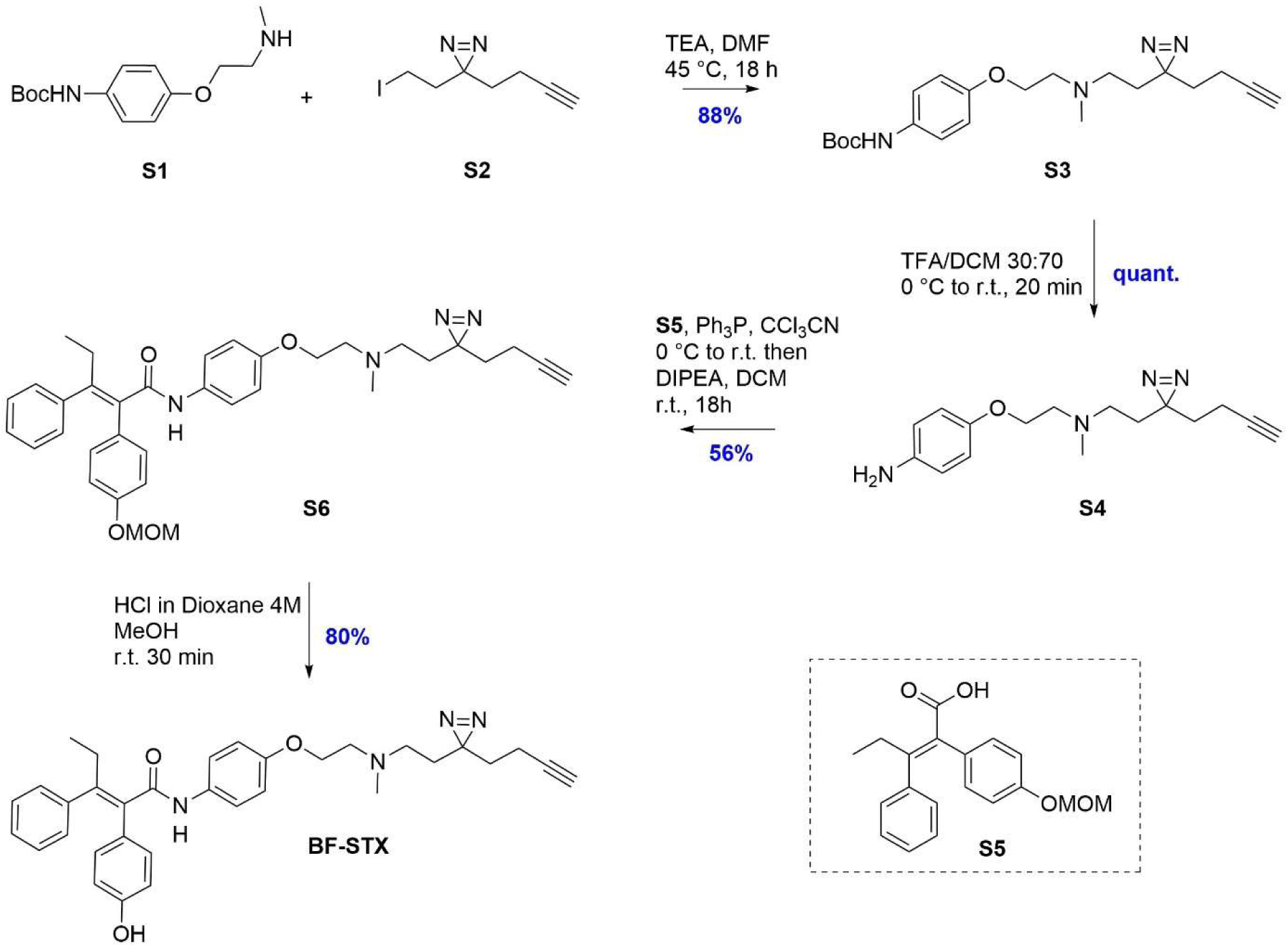
Synthesis scheme of BF-STX with reagents and yields.

**Figure S2.**
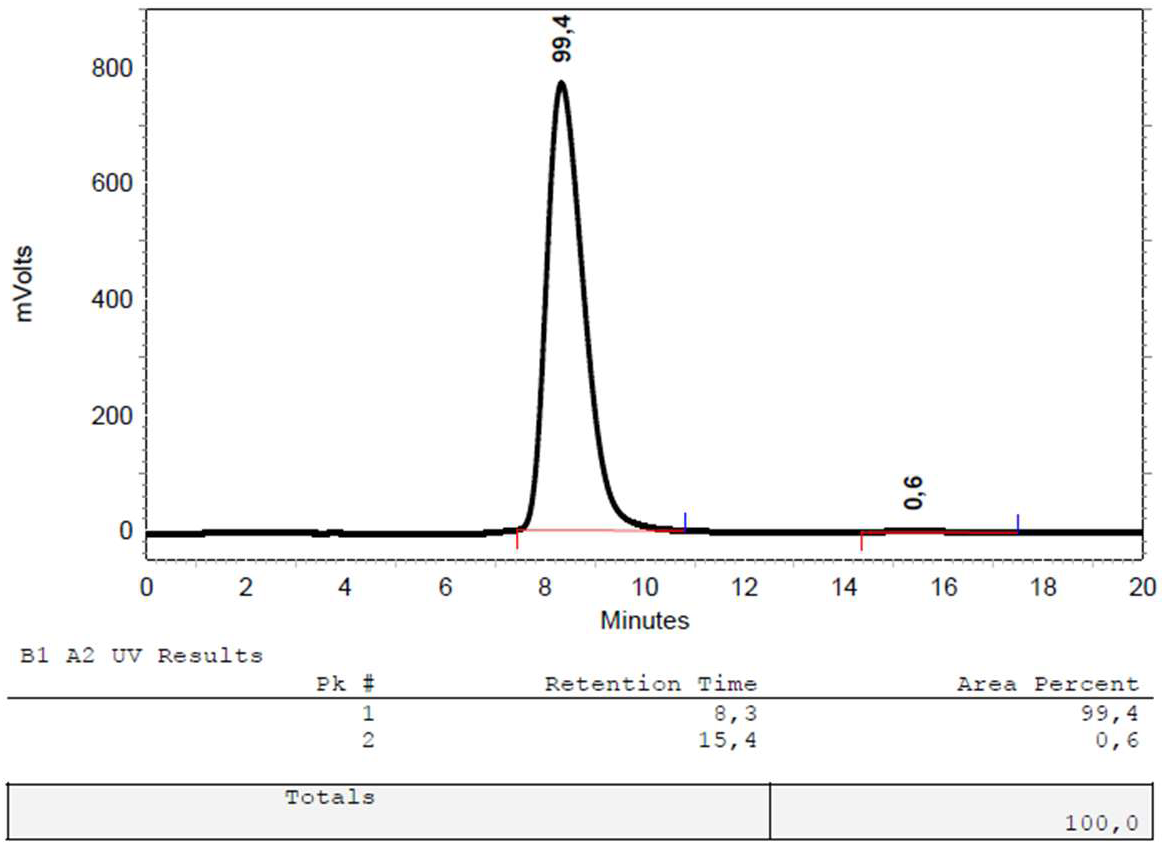
Analytical RP-HPLC chromatogram of final product BF-STX, according to Method A3.

**Figure S3.**
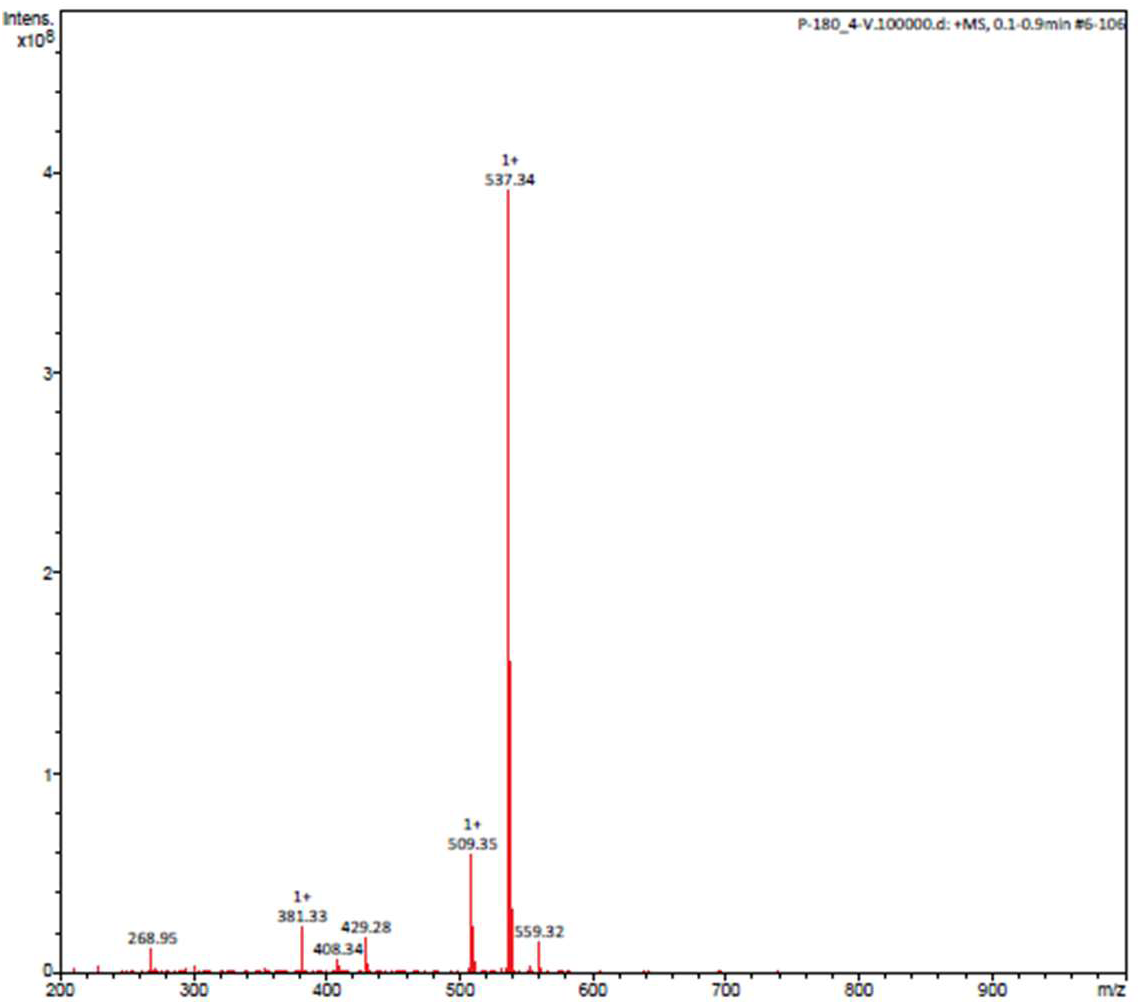
ESI-MS positive mode spectrum of final product BF-STX, showing m/z: 537.34 [M + H] ⁺ and m/z: 559.32 [M + Na] ⁺.

**Figure S4a.**
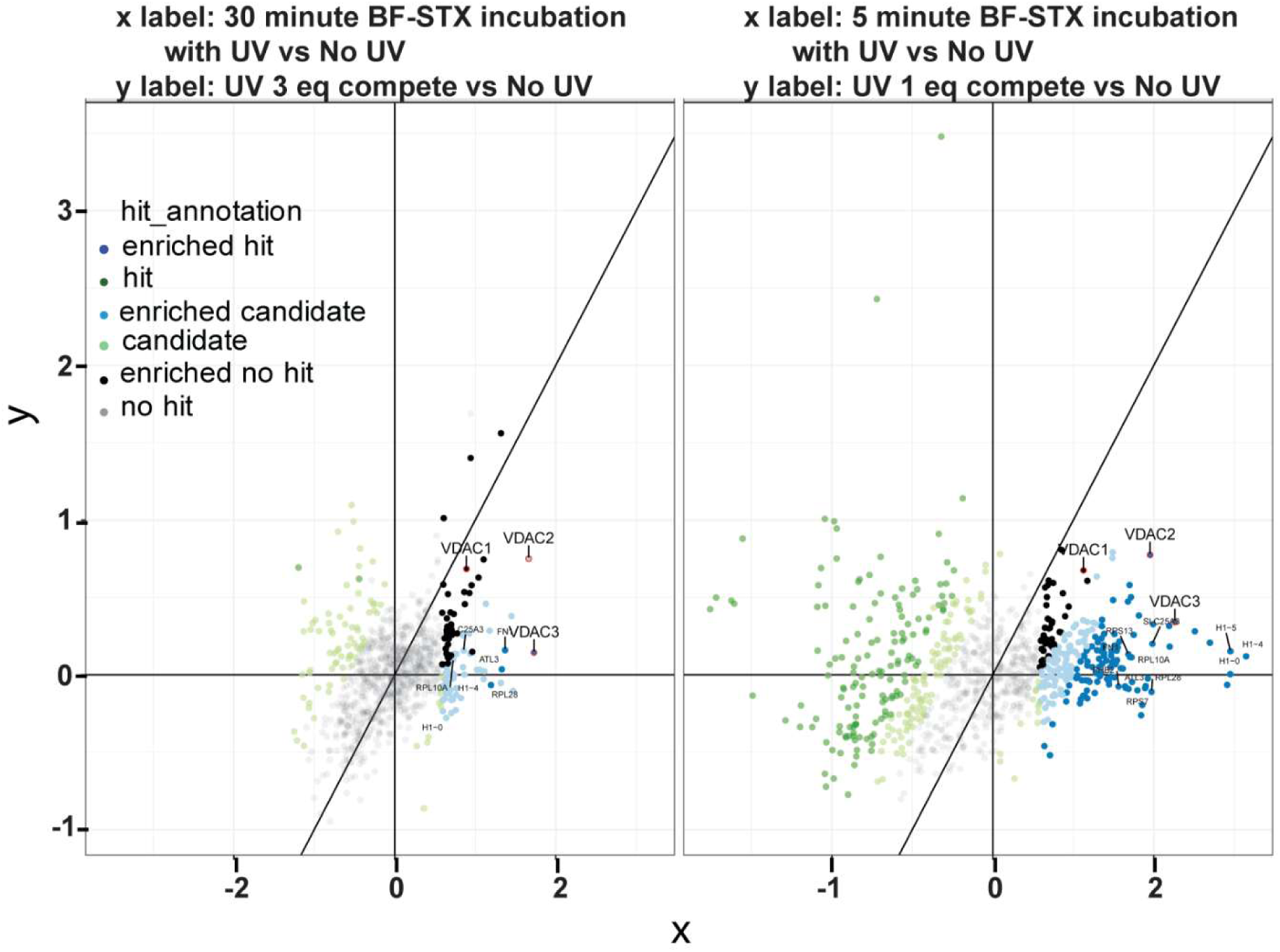
A. Fold-change correlation analysis of STX photoaffinity labeling and competition experiments. Scatter plots comparing protein enrichment fold changes under different STX photolabeling and competition conditions. In the left panel, the x-axis represents fold change following 30 min STX incubation with UV exposure versus no UV control, while the y-axis represents fold change in the presence of 3-equivalent unlabeled STX competitor versus no UV control. In the right panel, the x-axis represents fold change following 5 min STX incubation with UV exposure versus no UV control, while the y-axis represents fold change in the presence of 1-equivalent unlabeled STX competitor versus no UV control. Proteins enriched by UV-dependent labeling are distributed along the positive x-axis, whereas proteins showing reduced enrichment in the presence of competitor are consistent with competition-sensitive BF-STX labeling. VDAC1, VDAC2, and VDAC3 displayed robust UV are highlighted as candidate targets identified across conditions. The diagonal line indicates equal fold change between compared conditions.

**Figure S4.**
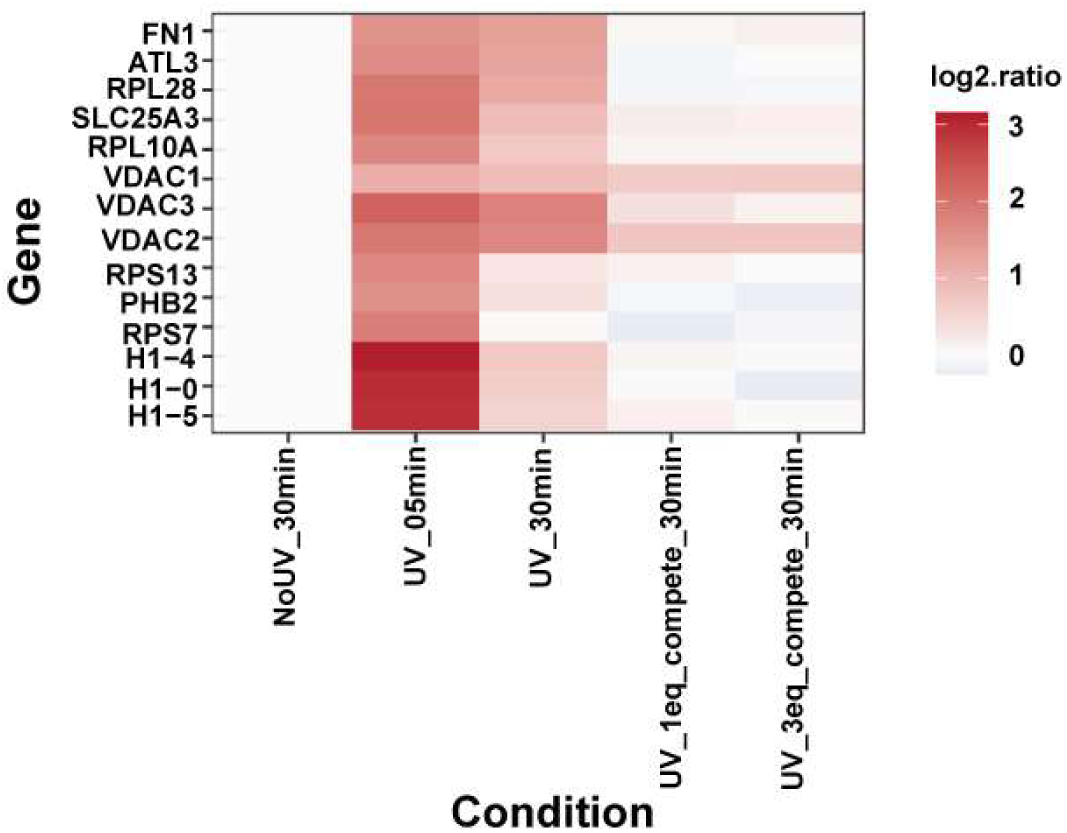
B. Heatmap analysis of candidate proteins identified in STX photoaffinity labeling experiments. Heatmap showing relative enrichment (log2 ratio) of candidate proteins detected across multiple STX photolabeling and competition conditions. Columns represent experimental conditions, including no UV control (NoUV_30min), UV crosslinking following 5 min or 30 min STX incubation (UV_05min and UV_30min), and UV crosslinking in the presence of 1-equivalent or 3-equivalent unlabeled STX competitor (UV_1eq_compete_30min and UV_3eq_compete_30min). Rows indicate identified candidate proteins. Color intensity corresponds to normalized log2 enrichment values, with darker red indicating greater enrichment. VDAC1, VDAC2, and VDAC3 displayed robust UV-dependent enrichment with partial reduction under competitive conditions, consistent with competition-sensitive BF-STX labeling. Histone proteins (H1-4, H1-0, and H1-5) and several ribosomal or mitochondrial-associated proteins were also detected with varying enrichment patterns across conditions.

**Figure S5.**
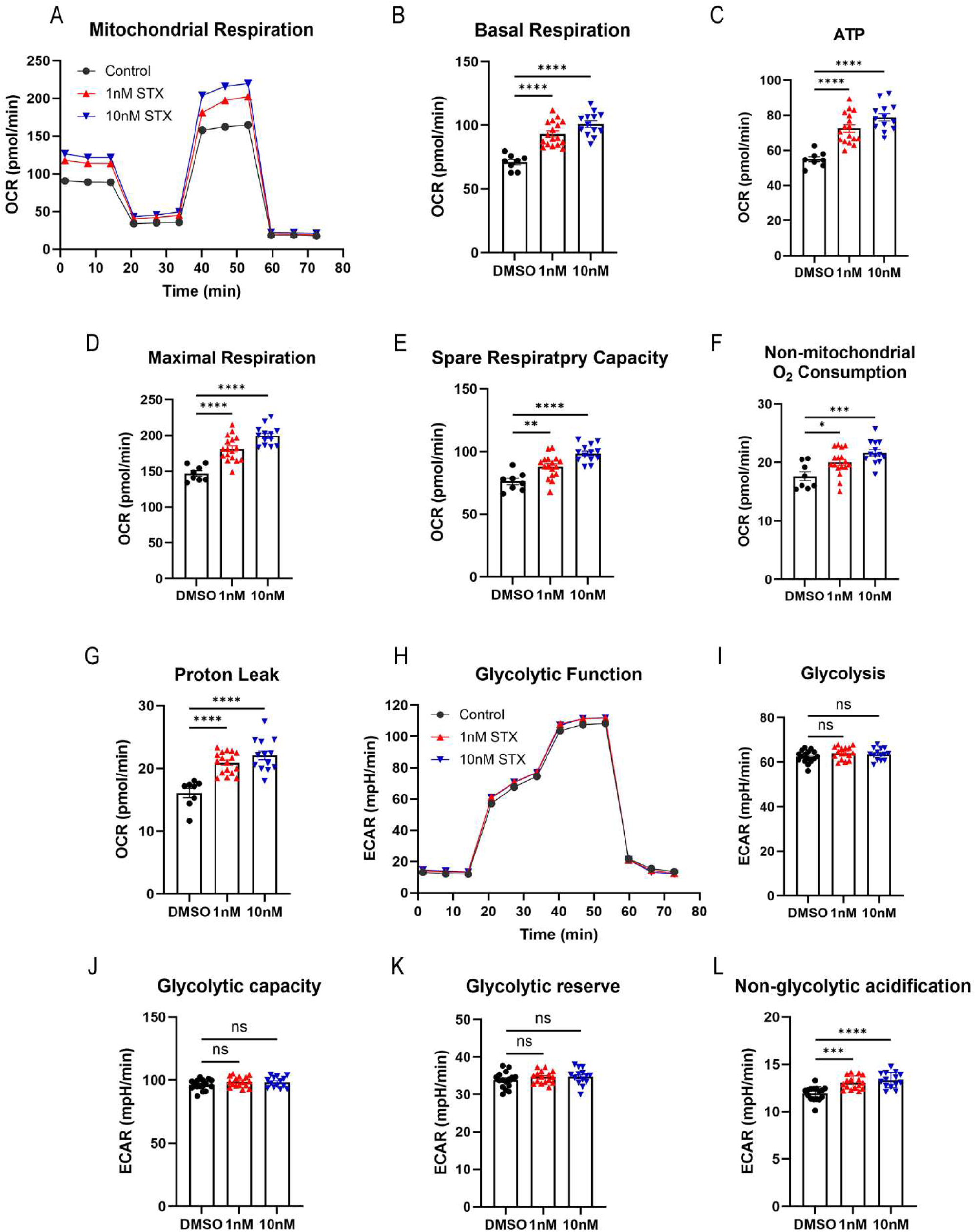
STX enhances mitochondrial respiration with minimal effects on glycolytic function in non–serum starved immortalized human hepatocytes (IHH). (A) Representative Seahorse XF traces of oxygen consumption rate (OCR) in non–serum starved immortalized human hepatocytes (IHH) treated with DMSO (control) or STX (1 nM, 10 nM). Sequential injections of oligomycin, FCCP, and rotenone/antimycin A were used to assess mitochondrial respiratory function. (B–G) Quantification of mitochondrial respiration parameters derived from OCR measurements, including basal respiration (B), ATP-linked respiration (C), maximal respiration (D), spare respiratory capacity (E), non-mitochondrial oxygen consumption (F), and proton leak (G). STX treatment significantly increased basal respiration, ATP-linked respiration, maximal respiration, spare respiratory capacity, proton leak and non-mitochondrial oxygen consumption at both 1 nM and 10 nM compared to control. (H) Representative extracellular acidification rate (ECAR) traces during the glycolysis stress test under the same treatment conditions. Sequential injections of glucose, oligomycin, and 2-deoxyglucose (2-DG) were used to assess glycolytic function. (I–L) Quantification of glycolytic parameters derived from ECAR measurements, including glycolysis (I), glycolytic capacity (J), glycolytic reserve (K), and non-glycolytic acidification (L). STX treatment did not significantly alter glycolysis, glycolytic capacity, or glycolytic reserve, but significantly increased non-glycolytic acidification. Data are presented as mean ± SEM with individual data points shown. Statistical significance was determined by one-way ANOVA followed by Dunnett’s multiple comparison test. ns, not significant; *p < 0.05, **p < 0.01, ***p < 0.001, ****p < 0.0001.

**Figure S6.**
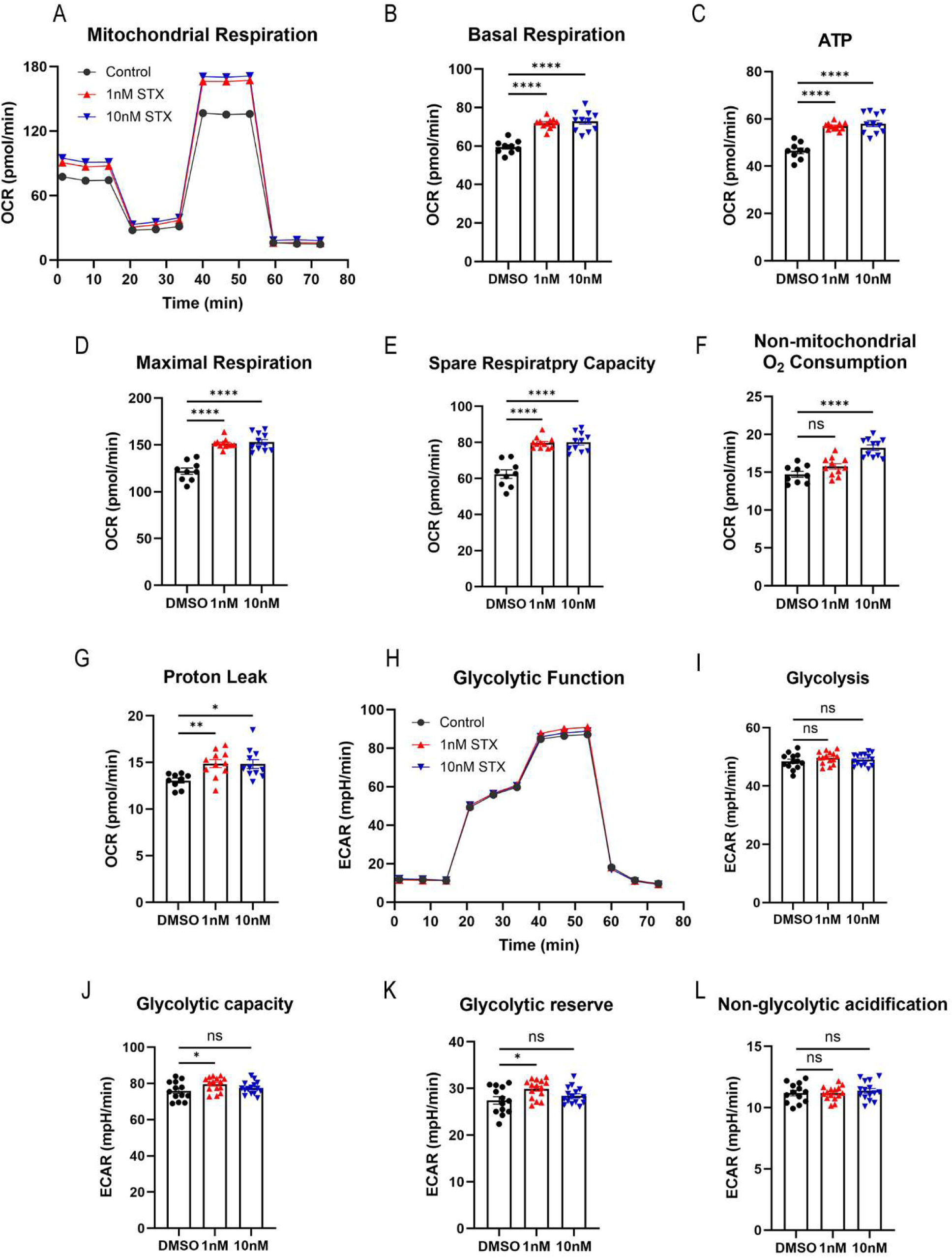
STX enhances mitochondrial respiration with minimal effects on glycolytic function in 6-hr serum starved IHH cells. (**A**) Representative Seahorse XF traces of oxygen consumption rate (OCR) in 6-hr serum starved immortalized human hepatocytes (IHH) treated with DMSO (control) or STX (1 nM, 10 nM). Sequential injections of oligomycin, FCCP, and rotenone/antimycin A were used to assess mitochondrial respiratory function. (**B–G**) Quantification of mitochondrial respiration parameters derived from OCR measurements, including basal respiration (**B**), ATP-linked respiration (**C**), maximal respiration (**D**), spare respiratory capacity (**E**), non-mitochondrial oxygen consumption (**F**), and proton leak (**G**). STX treatment significantly increased basal respiration, ATP-linked respiration, maximal respiration, and spare respiratory capacity at both 1 nM and 10 nM compared to control. Proton leak was also increased following the STX treatment, whereas non-mitochondrial oxygen consumption was significantly increased at 10 nM STX but not at 1 nM STX. (H) Representative extracellular acidification rate (ECAR) traces during the glycolysis stress test under the same treatment conditions. Sequential injections of glucose, oligomycin, and 2-deoxyglucose (2-DG) were used to assess glycolytic function. (**I–L**) Quantification of glycolytic parameters derived from ECAR measurements, including glycolysis (**I**), glycolytic capacity (**J**), glycolytic reserve (**K**), and non-glycolytic acidification (**L**). STX treatment did not significantly alter glycolysis, or non-glycolytic acidification. A modest increase in glycolytic capacity and glycolytic reserve was observed at 1 nM, whereas 10 nM STX did not significantly affect these parameters. Data are presented as mean ± SEM with individual data points shown. Statistical significance was determined by one-way ANOVA followed by Dunnett’s multiple comparison test. ns, not significant; *p < 0.05, **p < 0.01, ***p < 0.001, ****p < 0.0001.

## IV) NMR Spectra

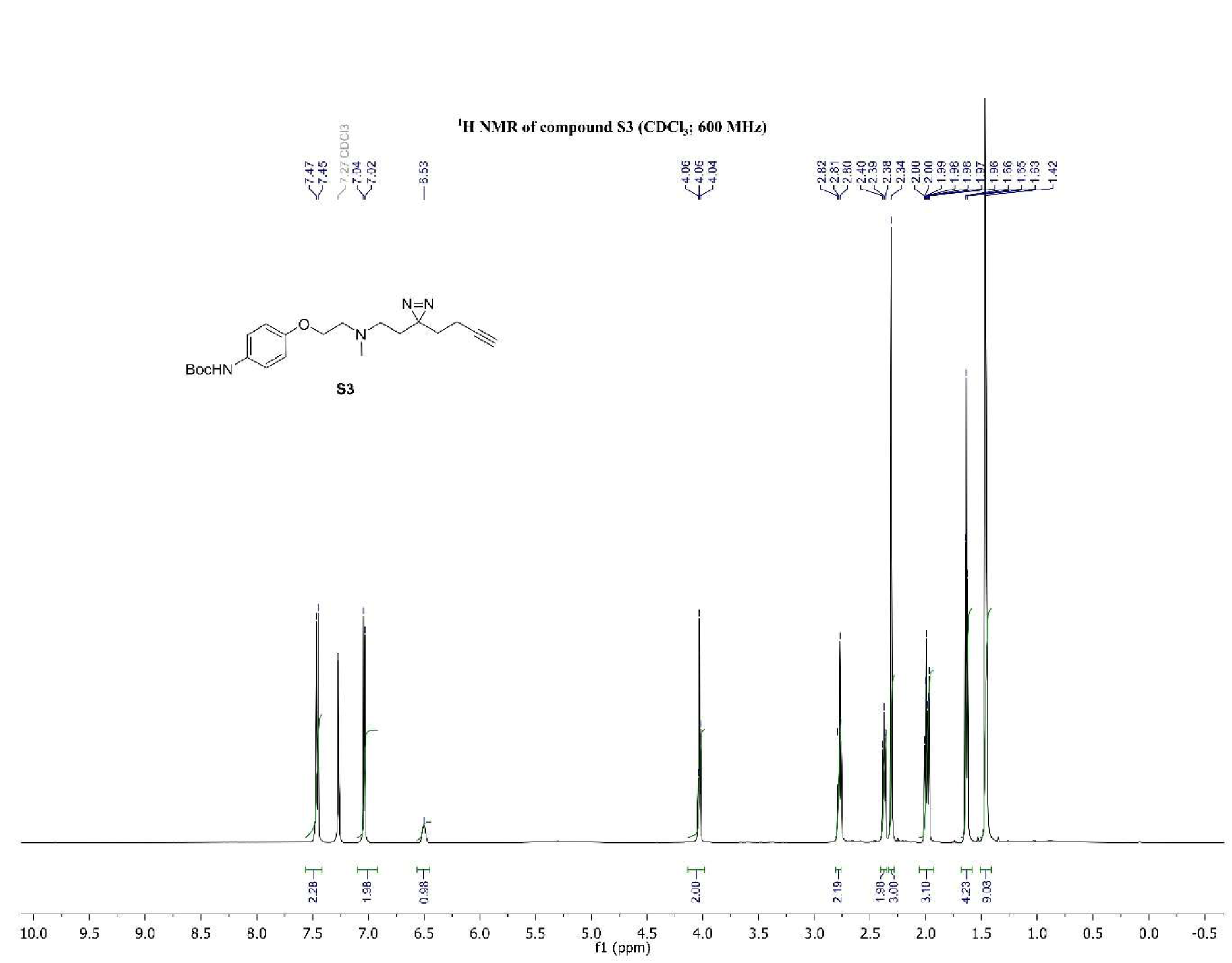
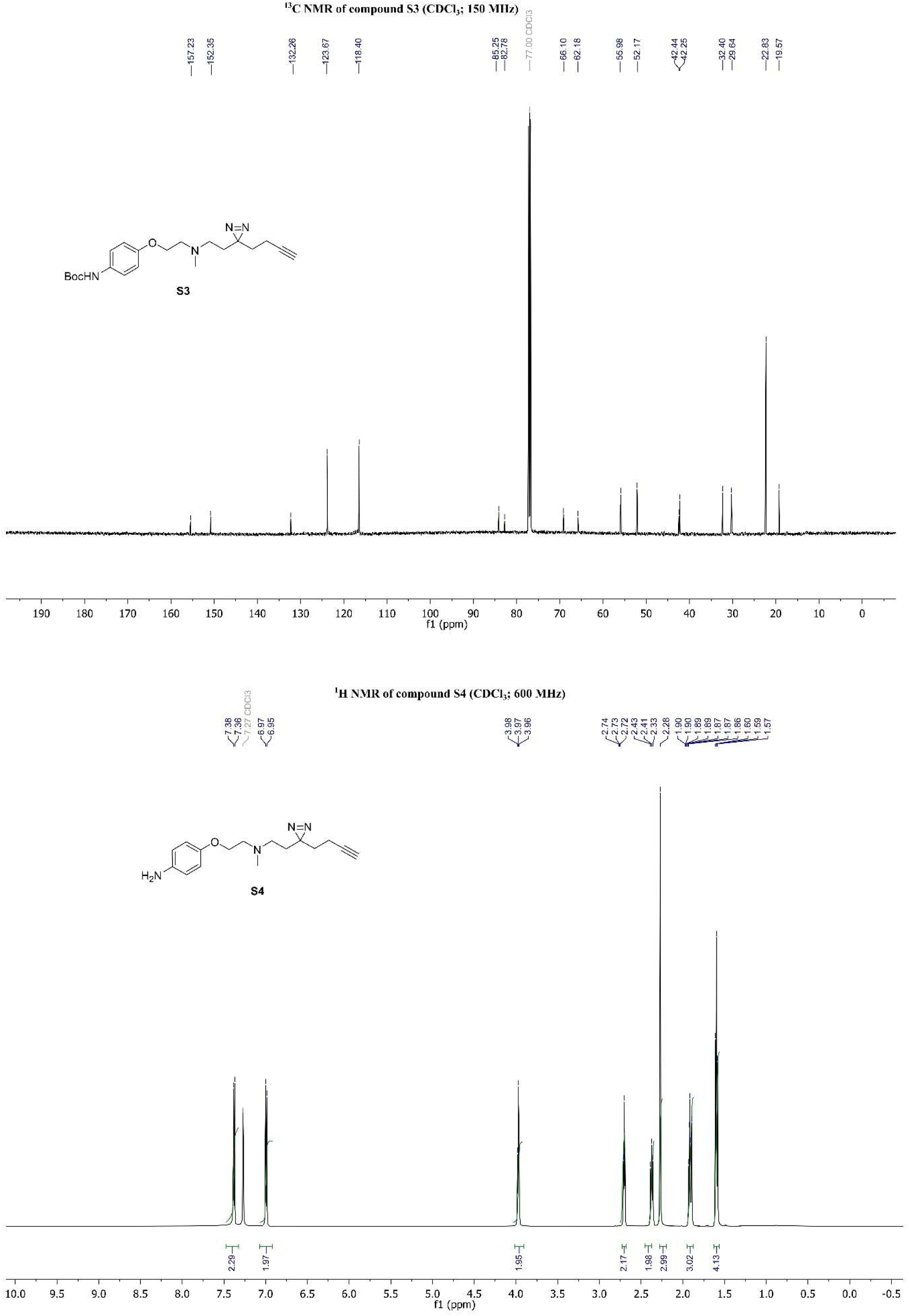
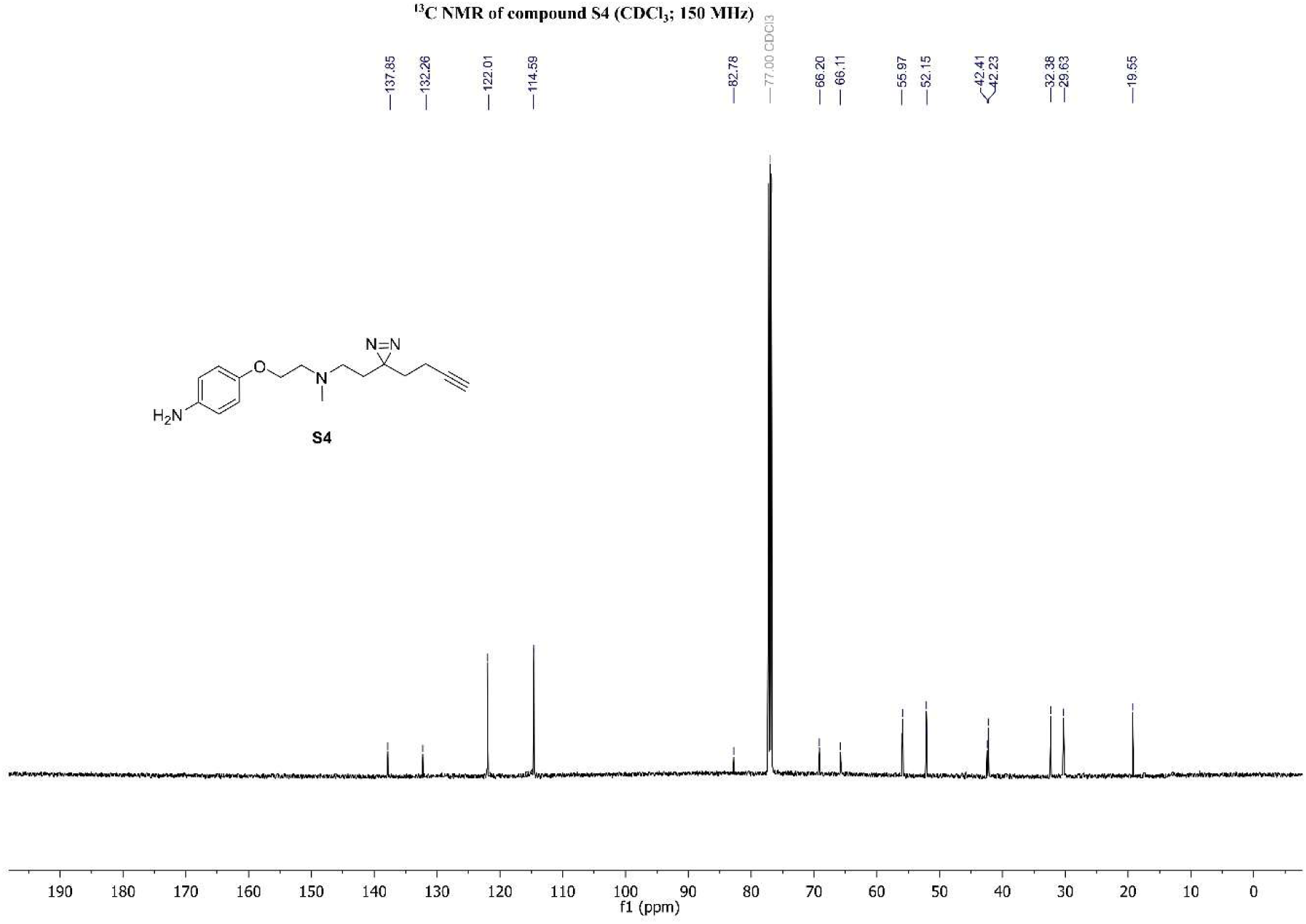
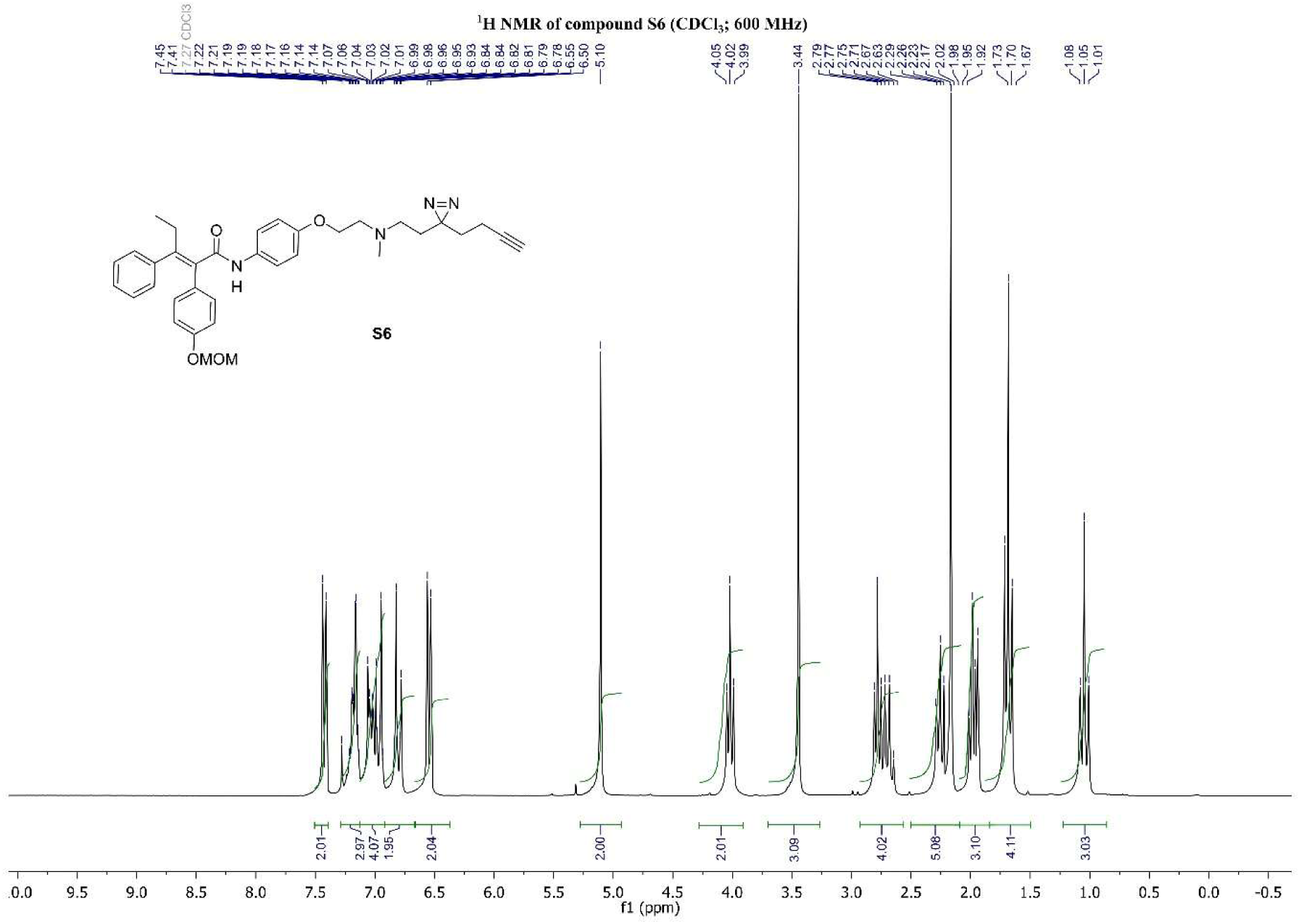
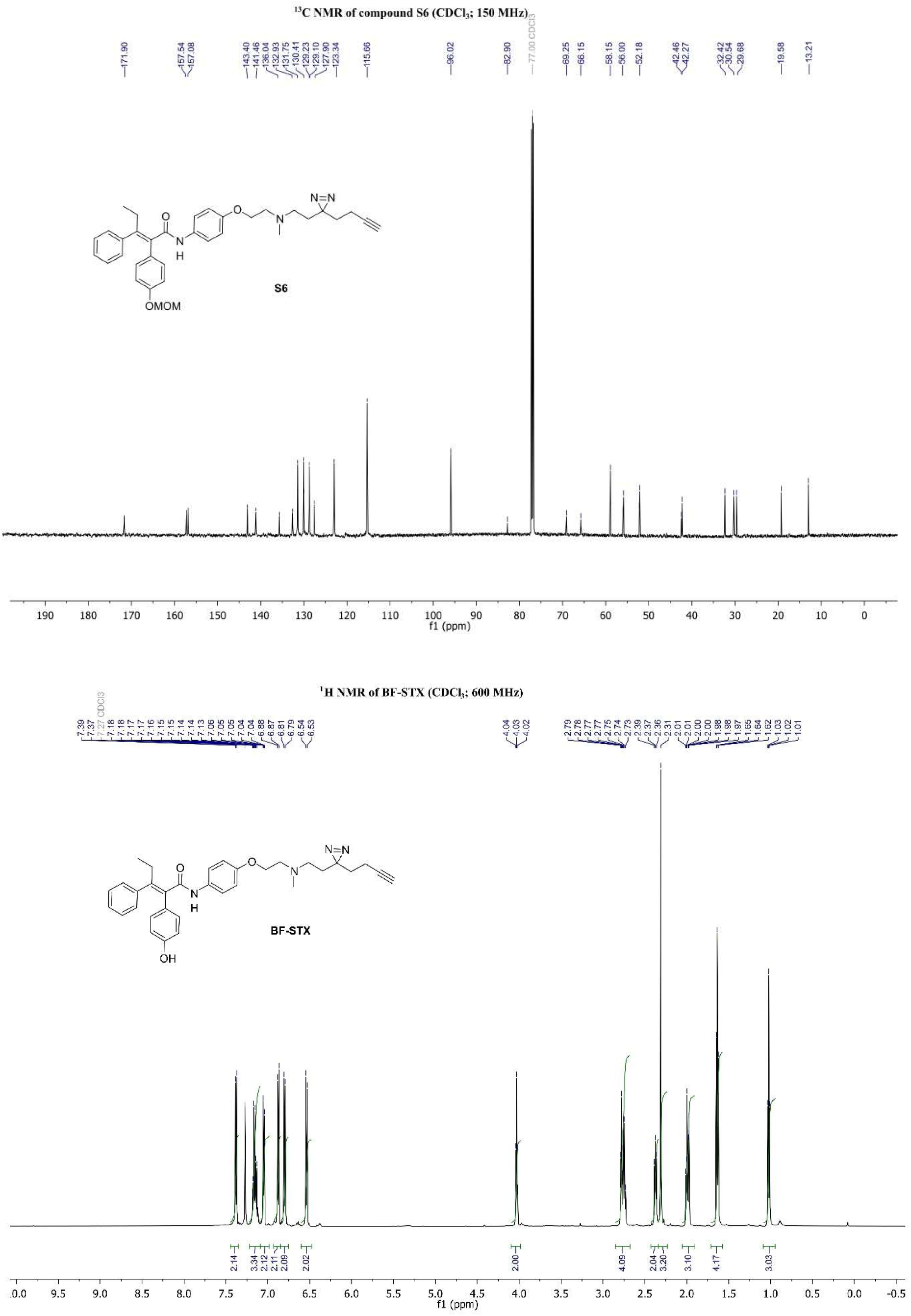
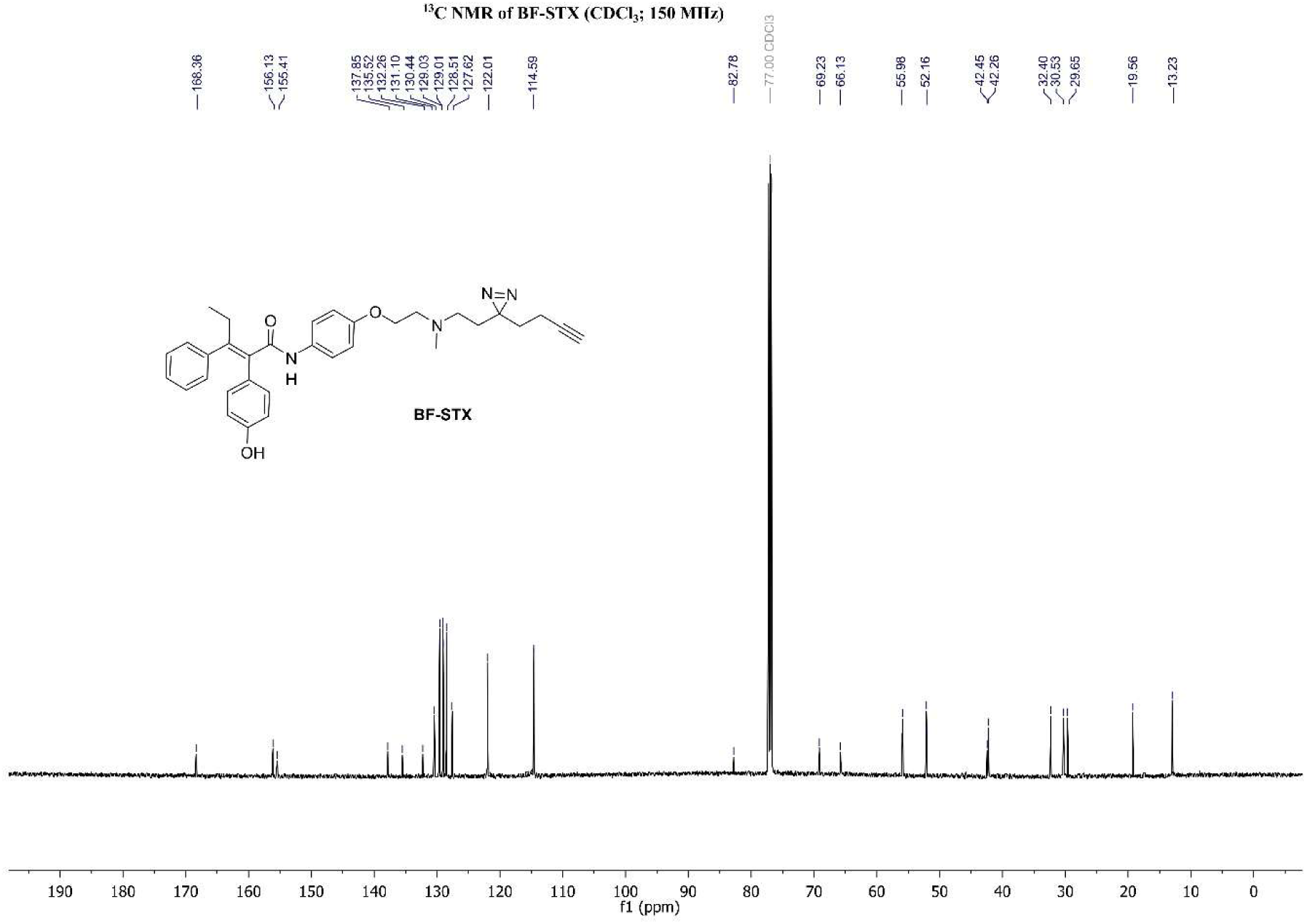

## V) Identification of isolated proteins by LC-MS/MS

At EMBL protein samples were subjected to the SP3 protocol ^1^ conducted on the KingFisher Apex™ platform (Thermo Fisher). For digestion, trypsin was used in a 1:20 ratio (protease:protein) in 50 mM N-2-hydroxyethylpiperazine-N-2-ethanesulfonic acid (HEPES) supplemented with 5 mM Tris(2-carboxyethyl)phosphine hydrochloride (TCEP) and 20 mM 2-chloroacetamide (CAA). Digestion was carried out for 5 hours at 37°C.

Up to 10 µg of peptides were labeled using TMTpro™ 16plex reagent as previously described ^2^. Briefly, 0.5 mg of TMT reagent was dissolved in 45 µL of 100% acetonitrile. Subsequently, 4 µL of this solution was added to each peptide sample, followed by incubation at room temperature for 1 hour. The labeling reaction was quenched by adding 4 µL of a 5% aqueous hydroxylamine solution and incubating for an additional 15 minutes at room temperature. Labeled samples were then combined for multiplexing, desalted using an Oasis® HLB µElution Plate (Waters) according to the manufacturer’s instructions, and dried by vacuum centrifugation.

An UltiMate 3000 RSLCnano LC system (Thermo Fisher Scientific) equipped with a trapping cartridge (µ-Precolumn C18 PepMap™ 100, 300 µm i.d. × 5 mm, 5 µm particle size, 100 Å pore size; Thermo Fisher Scientific) and an analytical column (nanoEase™ M/Z HSS T3, 75 µm i.d. × 250 mm, 1.8 µm particle size, 100 Å pore size; Waters) were used. Samples were trapped at a constant flow rate of 30 µL/min using 0.05% trifluoroacetic acid (TFA) in water for 6 minutes. After switching in-line with the analytical column, which was pre-equilibrated with solvent A (3% dimethyl sulfoxide [DMSO], 0.1% formic acid in water), the peptides were eluted at a constant flow rate of 0.3 µL/min using a gradient of increasing solvent B concentration (3% DMSO, 0.1% formic acid in acetonitrile). *The gradient was as follows: 2% to 8% in 4 minutes (min), 8% to 26% in 104 min, 26% to 38% in 4 min, 40%-80% in 0.1 min, 80% for 3.9 min and re-equilibrated to 2% B for 4 min*.

*Peptides were then introduced into an Orbitrap Fusion™ Lumos™ Tribrid™ mass spectrometer (Thermo Fisher Scientific) via a Pico-Tip emitter (360 µm OD × 20 µm ID; 10 µm tip, CoAnn Technologies) using an applied spray voltage of 2.2 kV. The capillary temperature was maintained at 275 °C. Full MS scans were acquired in profile mode over an m/z range of 375–1,500, with a resolution of 120,000 at m/z 200 in the Orbitrap. The maximum injection time was set to 50 ms, and the AGC target limit was set to ‘standard’. The instrument was operated in data-dependent acquisition (DDA) mode, with MS/MS scans acquired in the Orbitrap at a resolution of 30,000. The maximum injection time was set to 94 ms, with an AGC target of 200%. Fragmentation was performed using higher-energy collisional dissociation (HCD) with a normalized collision energy of 34%, and MS2 spectra were acquired in profile mode. The quadrupole isolation window was set to 0.7 m/z, and dynamic exclusion was enabled with a duration of 60 seconds. Only precursor ions with charge states 2–7 were selected for fragmentation*.

*Data analysis: Raw files were converted to mzML format using MSConvert from ProteoWizard, using peak picking, 64-bit encoding and zlib compression, and filtering for the 1000 most intense peaks. Files were then searched using MSFragger in FragPipe (version 19.0) against FASTA database UP000005640_HomoSapiens_ID9606_20594entries_26102022_dl11012023 containing common contaminants and reversed sequences. The following modifications were included into the search parameters: Carbamidomethylation (C, 57.0215), TMTpro (K, 304.2072) as fixed modifications; Oxidation (M, 15.9949), Acetylation (protein N-terminus, 42.0106), TMTpro (peptide N-terminus, 304.2072) as variable modifications. For the full scan (MS1) a mass error tolerance of 20 PPM and for MS/MS (MS2) spectra of 20 PPM was set. For protein digestion, ’trypsin’ was used as protease with an allowance of maximum 2 missed cleavages requiring a minimum peptide length of 7 amino acids. The false discovery rate on peptide and protein level was set to 0.01. The standard settings of the FragPipe workflow ’Default’ were used with several modifications: msfragger.isotope_error: -1/0/1/2/3, msfragger.misc.fragger.clear-mz-hi: 134.5, msfragger.misc.fragger.clear-mz-lo: 125.5, msfragger.misc.fragger.digest-mass-lo: 200, msfragger.misc.fragger.enzyme-dropdown-1: trypsin, msfragger.misc.fragger.precursor-charge-hi: 6, msfragger.search_enzyme_name_1: trypsin, msfragger.search_enzyme_nocut_1: P, msfragger.table.fix-mods.K.lysine.: 304.2072, msfragger.table.var-mods.peptide N-terminus.304.2072: peptide N-terminus, msfragger.use_topN_peaks: 300, phi-report.prot-level-summary: false, protein-prophet.cmd-opts: --maxppmdiff 2000000 --minprob 0.5, tmtintegrator.add_Ref: 1, tmtintegrator.channel_num: TMT-16, tmtintegrator.extraction_tool: Philosopher, tmtintegrator.max_pep_prob_thres: 0.9, tmtintegrator.run-tmtintegrator: true*.

*For the proteomics data analysis, the raw output files of FragPipe (protein.tsv files) were processed using the R programming environment (ISBN 3-900051-07-0). Initial data processing included filtering out contaminants and reverse proteins. Only proteins quantified with at least 2 unique peptides (Unique.Peptides >= 2) were considered for further analysis. 878 proteins passed the quality control filters. In order to correct for technical variability, batch effects were removed using the ’removeBatchEffect’ function of the limma package ^3^ on the log2 transformed raw TMT reporter ion intensities (’channel’ columns). Subsequently, normalization was performed using the ’normalizeVSN’ function of the limma package (VSN - variance stabilization normalization ^4^). For visualization, proteins abundances were normalized to the control condition by dividing by the median of control replicates (’ctrl’ column). These control ratios were used for clustering and heatmap display. Differential expression analysis was performed using the moderated t-test provided by the limma package ^3^. The model accounted for replicate information by including it as a factor in the design matrix passed to the ’lmFit’ function. To obtain p-values and false discovery rates (FDRs), the ’fdrtool’ function from the fdrtool package ^5^ was used to analyze the t-values produced by limma for certain comparisons. Proteins were annotated as enriched hits if they had a false discovery rate (FDR) below 0.05 and an absolute fold change greater than 2. Proteins were considered enriched candidates if they had an FDR below 0.2 and an absolute fold change greater than 1.5*.

---

**Supplemental Table 1.**
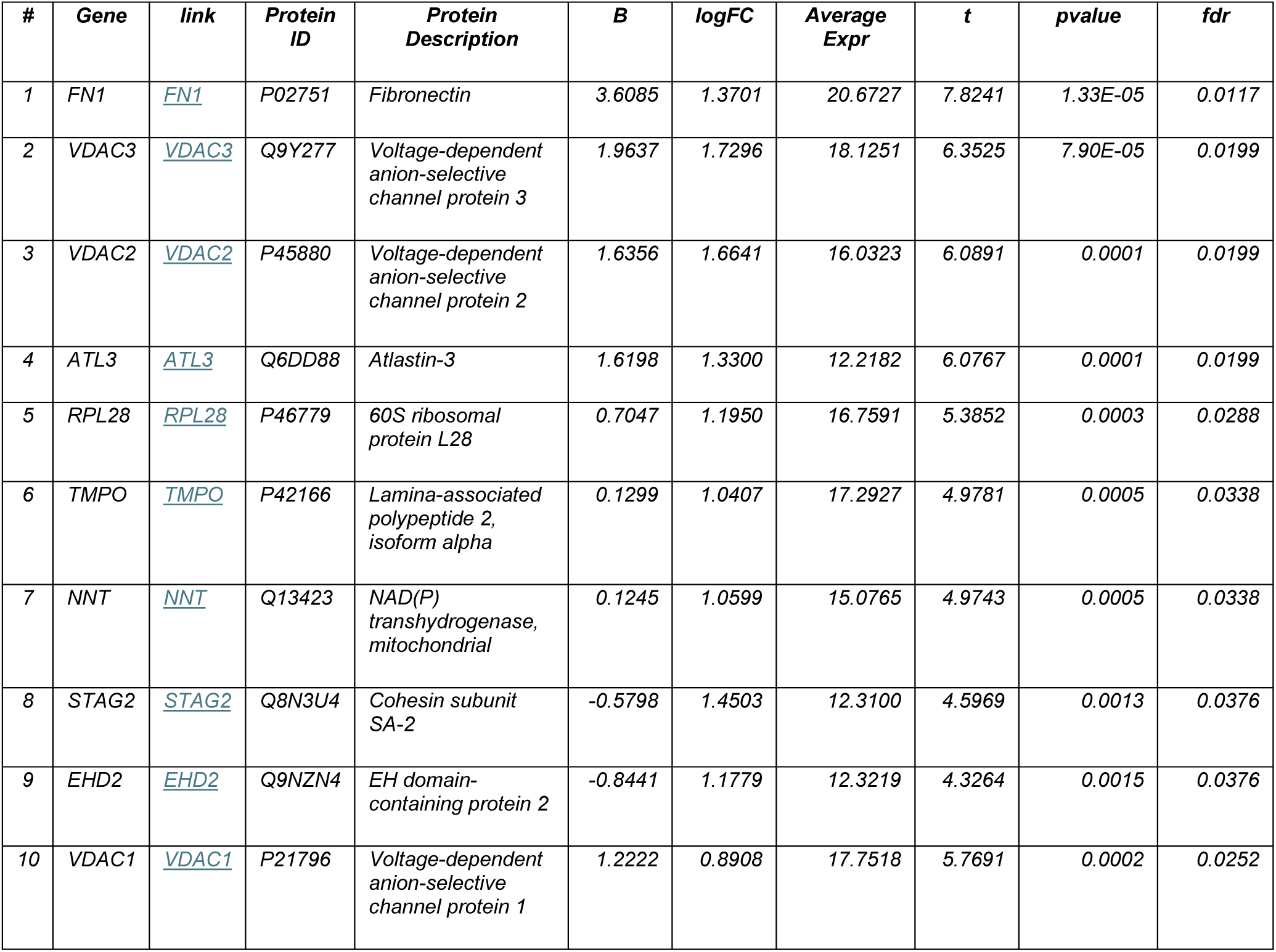
30 minute BF-STX incubation with UV minus No UV.

**Supplemental Table 2.**
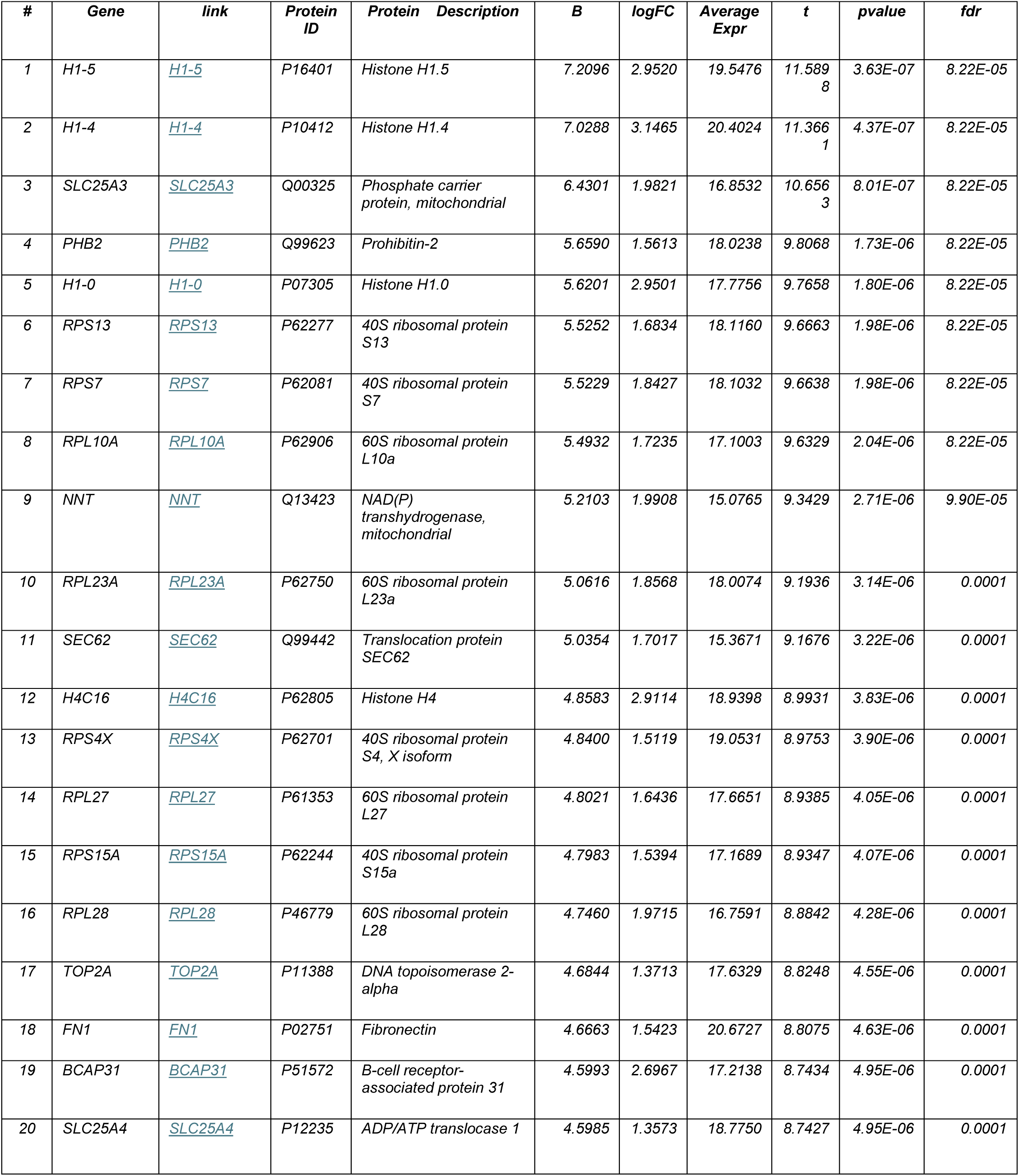
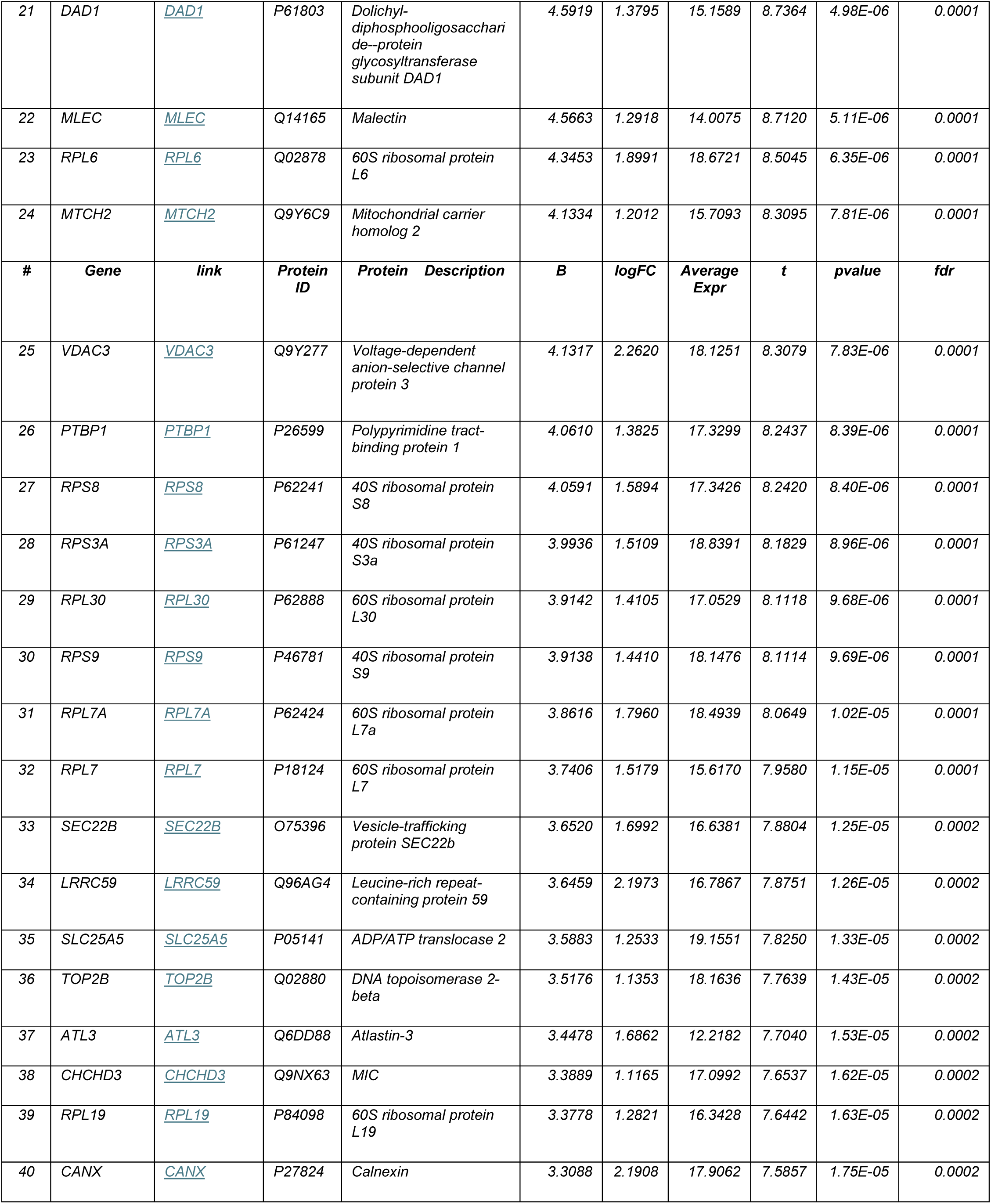
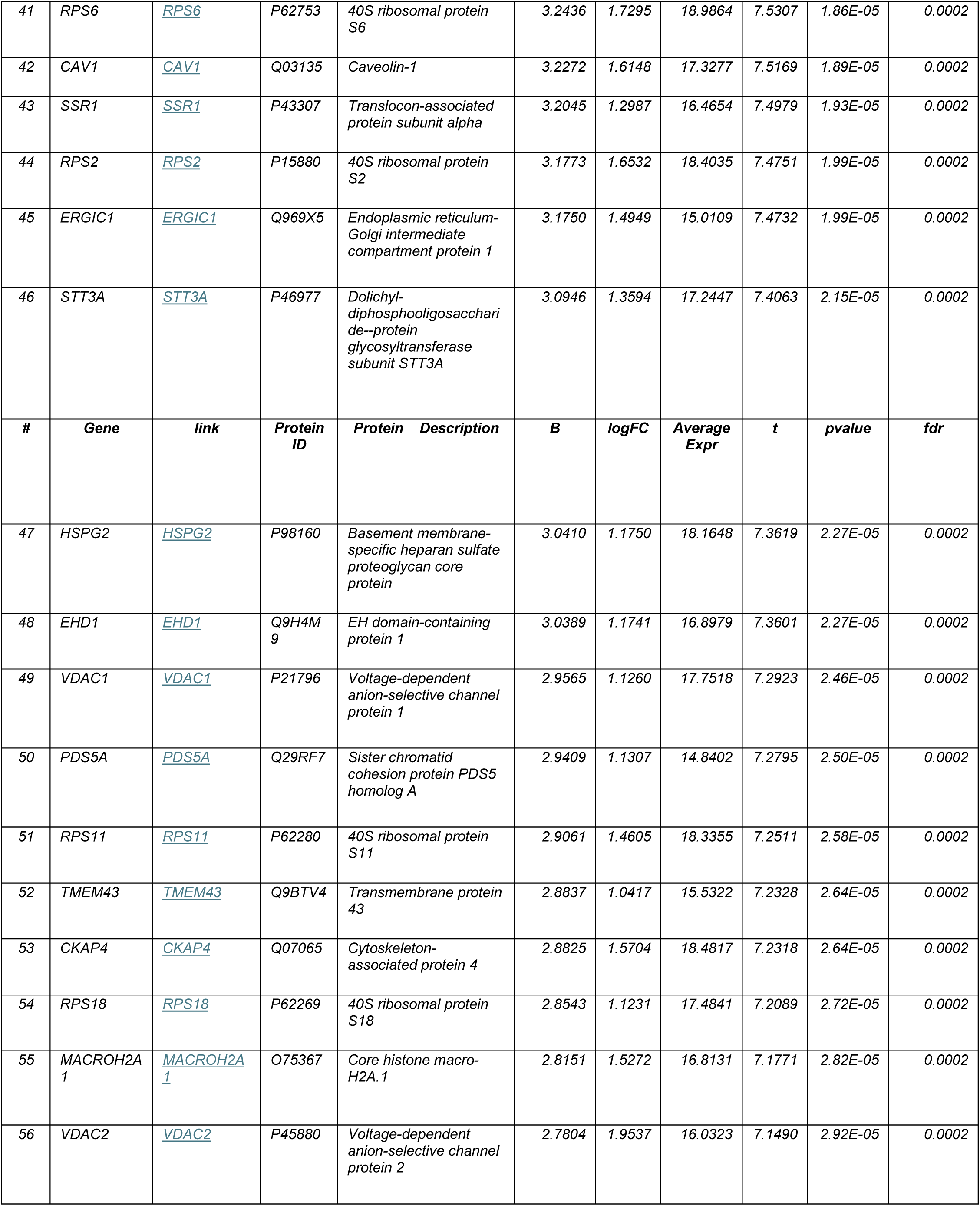
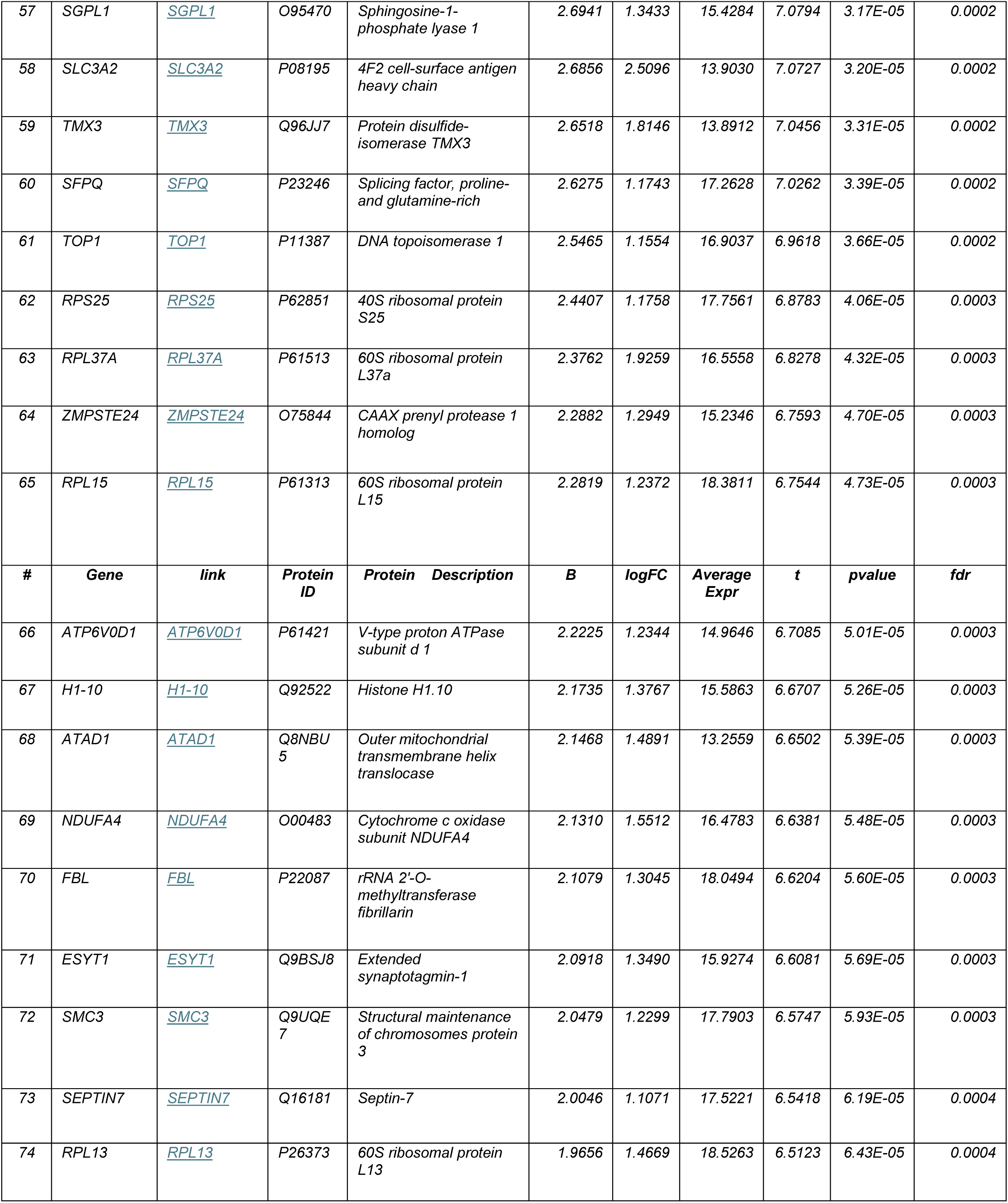
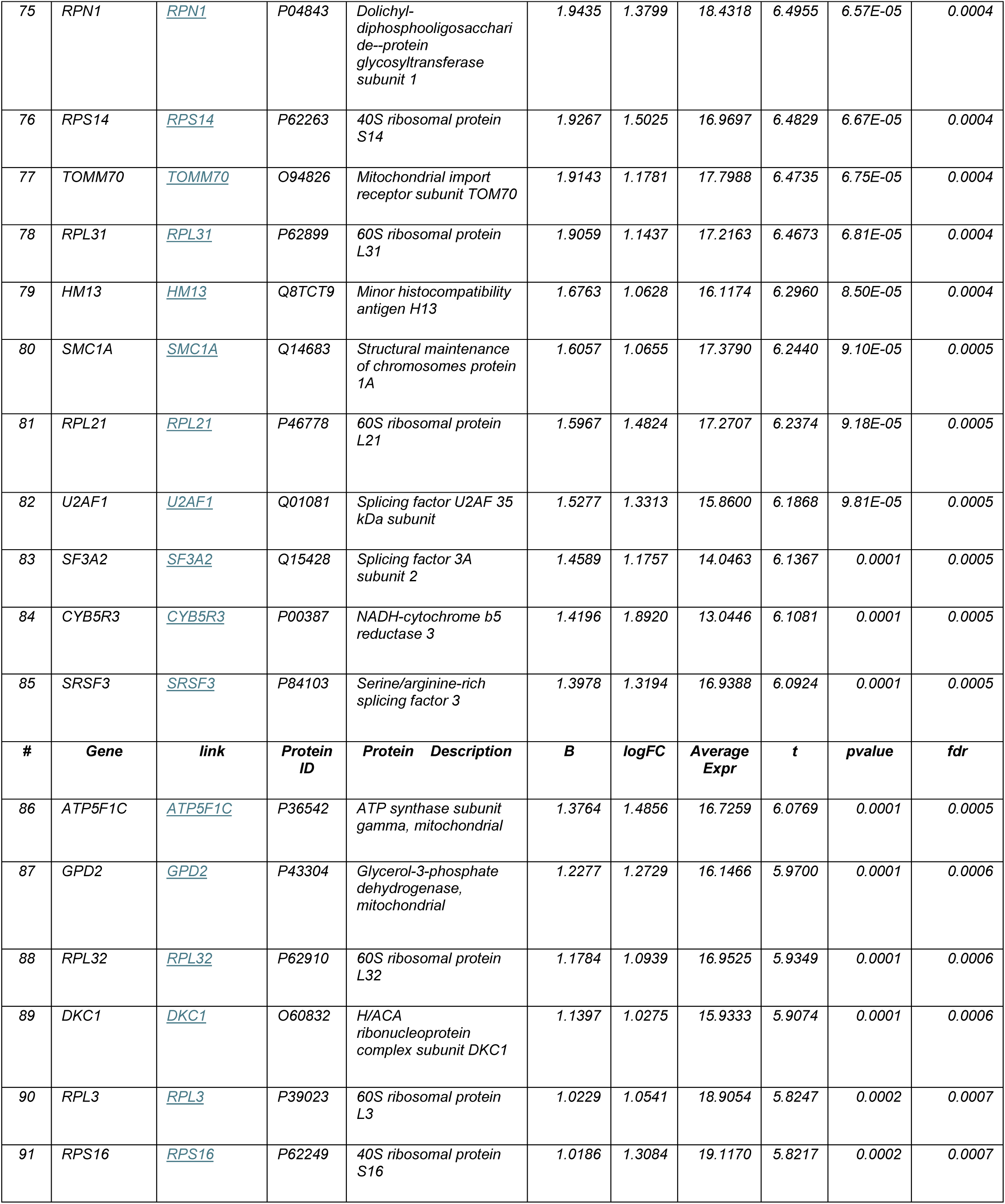
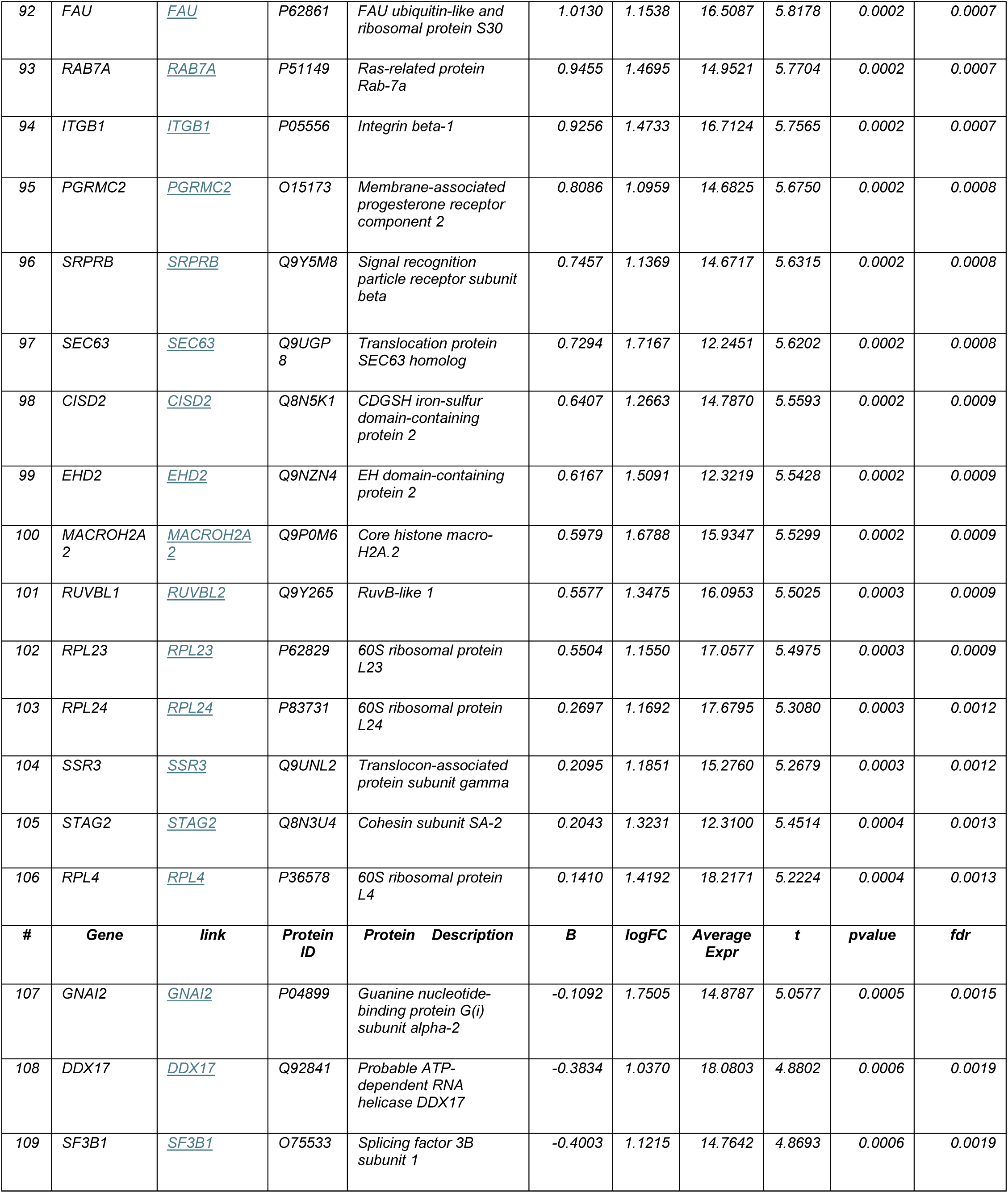
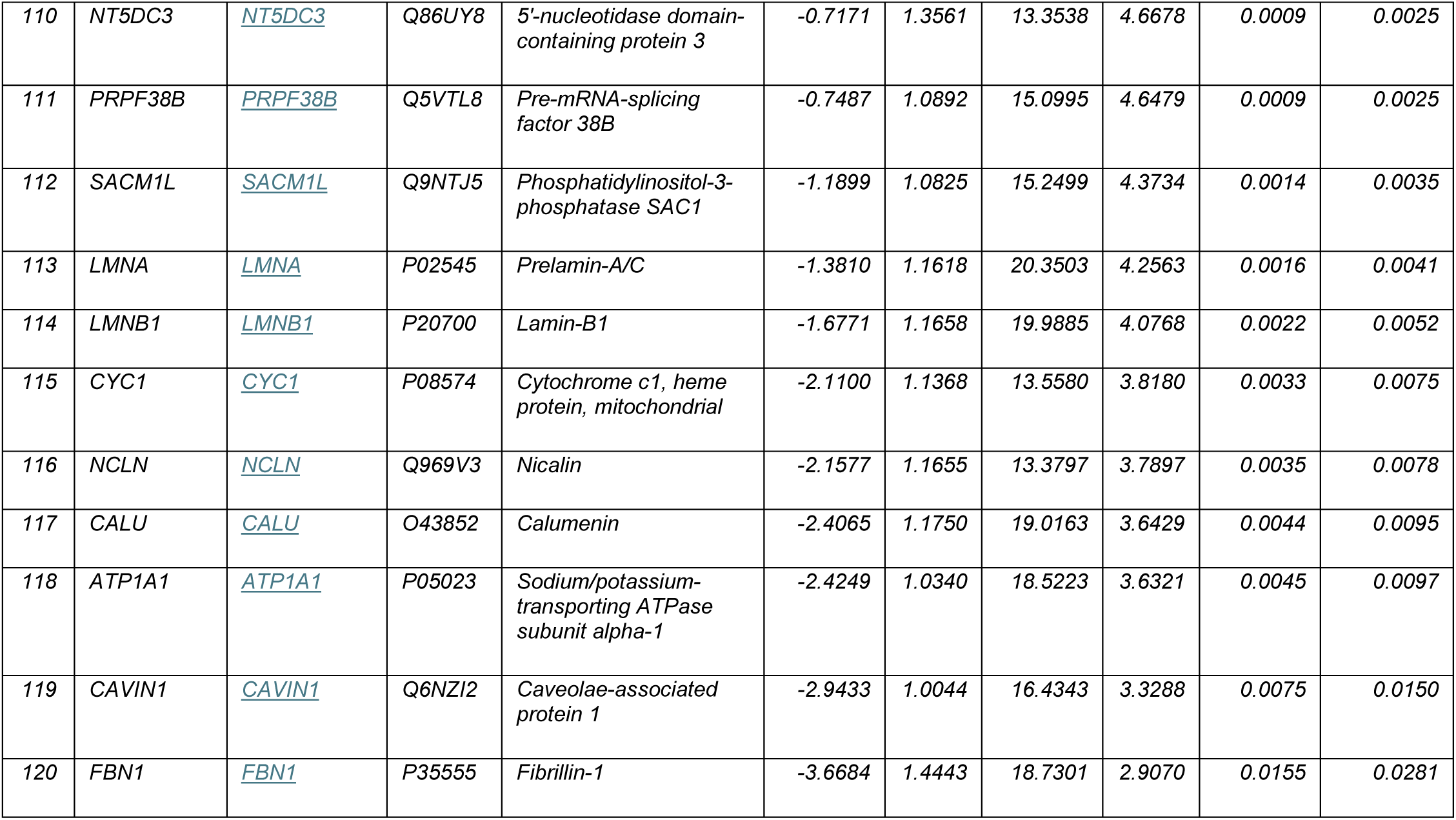
5 minute BF-STX incubation with UV minus No U.

